# Tandem paired nicking promotes precise genome editing with scarce interference by p53

**DOI:** 10.1101/810333

**Authors:** Toshinori Hyodo, Md Lutfur Rahman, Sivasundaram Karnan, Takuji Ito, Atsushi Toyoda, Akinobu Ota, Md Wahiduzzaman, Shinobu Tsuzuki, Yohei Okada, Yoshitaka Hosokawa, Hiroyuki Konishi

## Abstract

Targeted knock-in mediated by double-stranded DNA cleavage is accompanied by unwanted insertions and deletions (indels) at on-target and off-target sites. A nick-mediated approach scarcely generates indels but exhibits reduced efficiency of targeted knock-in. Here, we demonstrate that tandem paired nicking, a method for targeted knock-in involving two Cas9 nickases that create nicks at the homologous regions of the donor DNA and the genome in the same strand, scarcely creates indels at the edited genomic loci, while permitting the efficiency of targeted knock-in largely equivalent to that of the Cas9 nuclease-based approach. Tandem paired nicking seems to accomplish targeted knock-in via DNA recombination analogous to Holliday’s model, and creates intended genetic changes in the genome without introducing additional nucleotide changes such as silent mutations. Targeted knock-in through tandem paired nicking neither triggers significant p53 activation nor occurs preferentially in p53-suppressed cells. These properties of tandem paired nicking demonstrate its utility in precision genome engineering.

## Introduction

Genome editing using the CRISPR/Cas9 system is revolutionizing various research fields of life science (Cornu et al., 2017; Jinek et al., 2012; Komor et al., 2017). Targeted knock-in of foreign DNA fragments or base substitutions into genomes is believed to be one of particularly promising technologies, because this method permits modification of the genome exactly as designed. Targeted knock-in is generally conducted by introducing a DNA double-strand break (DSB) into the genome and then rendering it repaired by homology-directed repair (HDR) using an exogenous repair template called donor DNA. However, a majority of DSBs generated during genome editing are in fact repaired via non-homologous end-joining (NHEJ) rather than HDR, resulting in a high incidence of insertions and deletions (indels) at the DSB sites (Hsu et al., 2013; Mali et al., 2013). In addition, CRISPR/Cas9 often triggers unexpected cleavage followed by indel formation at off-target genomic sites (Fu et al., 2013; Hsu et al., 2013; Pattanayak et al., 2013; Ran et al., 2013a). Furthermore, it was recently shown that DSBs induced by CRISPR/Cas9 activate the p53 signaling pathway, and that wild-type p53 inhibits homology-directed genome engineering by CRISPR/Cas9 (Haapaniemi et al., 2018; Ihry et al., 2018). Technical improvements to overcome these issues, in addition to the maintenance of substantial efficiency of targeted knock-in, is a requisite to develop a method for precise and efficient genome engineering.

A genome editing approach named ‘base editing’ allows base substitution at an intended genomic site without introducing DSBs (Gaudelli et al., 2017; Komor et al., 2016; Nishida et al., 2016). Base editing is conducted using either a catalytically inactive Cas9 variant or a Cas9 nickase fused with an enzyme that deaminates cytidine or adenine. However, the current base editors can convert only C to T or A to G, may replace bystander Cs and As, does not insert or delete DNA, and edit only nucleotides located within an editing window near a protospacer-adjacent motif (PAM).

These caveats associated with DSB-mediated knock-in and base editing may be circumvented if efficient targeted knock-in via HDR can be achieved without creating DSBs in the genome. In this regard, previous studies have demonstrated that HDR is elicited by single-strand breaks (nicks) created in the genome using various site-specific nicking enzymes, including nick-only RAG mutants (Lee et al., 2004), a nicking variant of the I-AniI homing endonuclease (Davis and Maizels, 2011; McConnell Smith et al., 2009; Metzger et al., 2011), the adeno-associated virus (AAV) Rep protein (van Nierop et al., 2009), and Cas9 nickases (Cong et al., 2013; Mali et al., 2013). Although single nicks have been found to trigger HDR with efficiency generally lower than that triggered by DSBs (Bolukbasi et al., 2016; Cornu et al., 2017; Komor et al., 2017), further studies have shown that the efficiency of HDR can be improved by the use of two nickases so that they create no DSB in the genome. The majority of these studies created one nick on the donor DNA and the other within the genome, at different positions, using two nickases (Chen et al., 2017; Davis and Maizels, 2014; Goncalves et al., 2012; Nakajima et al., 2018). In this study, we explored a distinct and less well described strategy of using Cas9 nickases for targeted knock-in, which is called ‘tandem paired nicking’ (Chen et al., 2017), or ‘tandem nicking’ (Nakajima et al., 2018). In tandem paired nicking, two Cas9 nickases create nicks in the same strand, at homologous regions of the donor plasmid and the genome, in such a way that the nicks surround the knock-in site (Figure 1A). With this design, the nicks are placed at the same positions on the donor and the genome, and a total of four nicks are introduced by the two nickases. This nicking design was chosen because it permits targeted knock-in without additional nucleotide changes, such as silent mutations.

**Figure 1.**
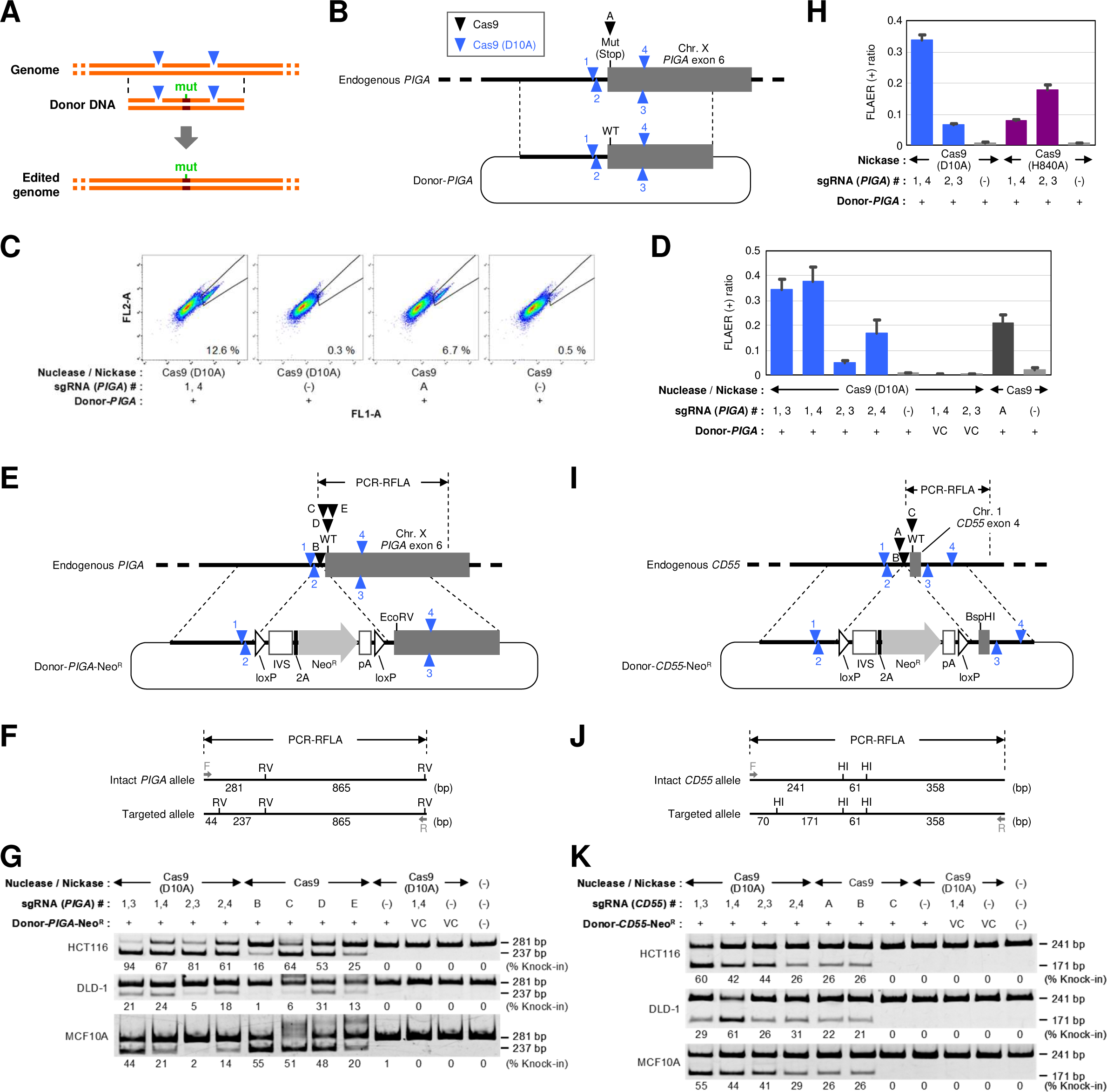
Tandem paired nicking promotes targeted knock-in with substantial efficiency. (A) Schematic description of tandem paired nicking. (B) Correction of a *PIGA* inactivating mutation via tandem paired nicking. An HCT116 clone carrying a *PIGA* mutation (HCT116-mut*PIGA*) was transfected with the Donor-*PIGA* plasmid along with a Cas9 nuclease or nickases, stained with FLAER, and analyzed by FCM (*PIGA* correction assay). Hereafter, schemes are not drawn to scale, and letters and numbers near arrowheads represent individual Cas9 nucleases and nickases in each target gene, respectively. Mut (Stop): an inactivating mutation constituting an ectopic stop codon. WT: a wild-type codon. (C) Representative dot plots obtained in FCM analyses for the *PIGA* correction assay. The percentage of FLAER-positive cells is noted within each plot. (D) Ratios of FLAER-positive cells representing the efficiencies of *PIGA* correction. Mean and s.e.m. values of biological triplicates are shown. (E) Targeted knock-in of a *PIGA* mutation via tandem paired nicking. Cell lines were transfected with Donor-*PIGA*-Neo^R^ along with a Cas9 nuclease or nickases, selected with G418, and processed for RFLA. EcoRV: a *PIGA*-inactivating mutation constituting an EcoRV site. This *PIGA* mutation is identical to the mutation depicted in (B). PCR-RFLA: a region amplified for RFLA. IVS: synthetic intron, 2A: ribosome skipping 2A sequence, Neo^R^: neomycin resistance (neomycin phosphotransferase) gene, pA: polyadenylation site. Recombination mediated by loxP was not carried out in this study. (F) Schematic description of RFLA for the targeted knock-in of a *PIGA* mutation. RV: EcoRV sites, F and R: PCR primers. (G) Results of RFLA for the targeted knock-in of a *PIGA* mutation. Percentages of knock-in alleles were calculated based on band intensities and are denoted under the corresponding lanes. (H) *PIGA* correction assay using Cas9 (D10A) and Cas9 (H840A). Mean and s.e.m. values of biological triplicates are shown. (I) –(K) Scheme (I and J) and results (K) of the targeted knock-in of a *CD55* mutation via tandem paired nicking, depicted as analogous to (E)–(G) for *PIGA*. BspHI and HI: BspHI sites. See also Figures S1–S3

We applied this method to edit multiple genetic loci in several human cell lines, and evaluated the frequencies of targeted knock-in and indel formation. In addition, we addressed whether this method triggers p53 activation, and whether functional p53 suppresses targeted knock-in accomplished with this method.

## Results

### Tandem paired nicking promotes targeted knock-in with substantial efficiency

We initially assessed the efficiency of nucleotide substitution via tandem paired nicking at endogenous genomic loci in human cell lines. Targeted knock-in via tandem paired nicking has been described previously; the efficiency of genome integration of an EGFP expression cassette was assessed in an assay based on the AAVS1 locus in the HeLa cell line (Chen et al., 2017), and the efficiency of nucleotide substitution via tandem nicking (tandem paired nicking) was assessed based on a stably transduced EGFP reporter cassette in 293T cells (Nakajima et al., 2018). Collectively, however, targeted knock-in via tandem paired nicking has been relatively less characterized than other nick-based knock-in strategies.

To delineate targeted knock-in via tandem paired nicking in distinct experimental settings, we initially generated a cell clone derived from the HCT116 cell line in which an ectopic stop codon is incorporated within exon 6 of the *PIGA* gene (hereafter called the HCT116-mut*PIGA* clone). *PIGA*, located on the X chromosome (Takeda et al., 1993), is inactivated in this clone, as HCT116 is a male cell line and has a single *PIGA* allele. Correction of the *PIGA* mutation in this clone is readily detectable by staining of the cells with an Alexa 488-conjugated inactive aerolysin variant (FLAER), a fluorescent reagent that binds specifically to glycosylphosphatidylinositol (GPI)-anchors on the cell membrane, followed by flow cytometric analysis (Brodsky et al., 2000; Miyata et al., 1993). This clone was initially transfected with Cas9 (D10A) (Jinek et al., 2012), a Cas9 variant that nicks the targeted strand, along with two single guide RNAs (sgRNAs) and a plasmid harboring a donor fragment that corrects the *PIGA* mutation (Figure 1B). We found that a subset (5–38 %) of cells became FLAER-positive after nicking at two distinct positions on the same DNA strand, in addition to ‘paired nicking’ in which two nicks are introduced into complementary strands (Figures 1C and 1D). This result suggests that a process distinct from DSB repair is involved in this gene correction. Subsequently, a *PIGA* mutation was introduced into parental HCT116 cells using the same set of Cas9 nickases along with a donor plasmid carrying a neomycin resistance (Neo^R^) gene (Figure 1E). Restriction fragment length analysis (RFLA) indicated that the knock-in efficiency using Cas9 nickases under these experimental conditions reached 90 % after enrichment based on G418 selection (Figures 1F and 1G).

In both of the above assays, tandem paired nicking stimulated targeted knock-in to achieve efficiencies roughly comparable to those of the Cas9 nuclease-based method. Similar results were obtained when the DLD-1 and MCF10A cell lines were employed in the RFLA-based assay detecting the targeted knock-in of a *PIGA* mutation (Figure 1G). In addition to Cas9 (D10A), the use of Cas9 (H840A) (Jinek et al., 2012), a Cas9 variant that nicks a non-targeted strand, also resulted in robust targeted knock-in via tandem paired nicking, as demonstrated by the *PIGA* correction assay using the HCT116-mut*PIGA* clone (Figure 1H). With one-base pair (bp) substitution within exon 4 of *CD55* (chromosome 1) followed by G418-based enrichment, tandem paired nicking again permitted targeted knock-in with efficiencies equivalent to those of Cas9 nucleases (Figures 1I–1K). In this analysis, one of three Cas9 nucleases (C) did not appreciably trigger targeted knock-in for unknown reasons, but demonstrated robust nuclease activity in the subsequent experiment (Figure S1F).

To confirm that no significant DSB is introduced via tandem paired nicking, we next performed an Indel Detection by Amplicon Analysis (IDAA) assay, in which size alteration of polymerase chain reaction (PCR) products indicates indel formation resulting from NHEJ repair of DSBs (Yang et al., 2015). We detected no appreciable size-altered PCR products in the cells subjected to tandem paired nicking using either Cas9 (D10A) or Cas9 (H840A), while cell transfection with Cas9 nucleases resulted in overt size alteration of the PCR products, indicating indel formation (Figure S1). Paired nicking with the PAM-out configuration led to a minor fraction of PCR products with size alterations upon *PIGA* editing but not *CD55* editing, probably because the distance between two complementary nicks in *PIGA* is shorter than that in *CD55* (261 nt versus 366 nt).

We also performed a so-called EGxxFP assay, in which GFP signal indicates the occurrence of a DSB in a transiently transfected reporter plasmid (Mashiko et al., 2013) (Figure S2A). As expected, no appreciable GFP signal was detected by cleavage of reporters with single nickases. However, complementary nicking of the reporters (with 13–247 nt overhangs) resulted in fair to robust GFP emission from cells, indicating that these Cas9 nickases indeed bind to and nick the transfected plasmids (Figure S2B).

We additionally co-cultured *PIGA*- or CD55-disrupted cells with their wild-type counterparts (Figure S3). No significant changes in the proportions of FLAER and anti-CD55 negative cells were observed over time. Similar to our previous results for *PIGA* (Karnan et al., 2012), these data validate the measurement of *PIGA*- and *CD55*-disruption ratios after a cell propagation period as indicators of gene editing efficiency. Collectively, these data indicate that tandem paired nicking achieves the efficiency of targeted knock-in largely equivalent to that of Cas9 nuclease-based approach, in line with previous results obtained in HeLa (Chen et al., 2017).

### Tandem paired nicking leads to DNA recombination analogous to Holliday’s model

Tandem paired nicking is reminiscent of the initial step in Holliday’s model, which was originally proposed to explain the genetic recombination that occurs during meiosis in fungi (Figure 2A) (Holliday, 1964; Liu and West, 2004). Holliday’s model proposed that isolocal nicking in two chromatids leads to the formation of a Holliday junction, followed by its resolution into either a crossover or non-crossover structure. As the branch in a Holliday junction migrates, gene conversion can accompany recombination.

**Figure 2.**
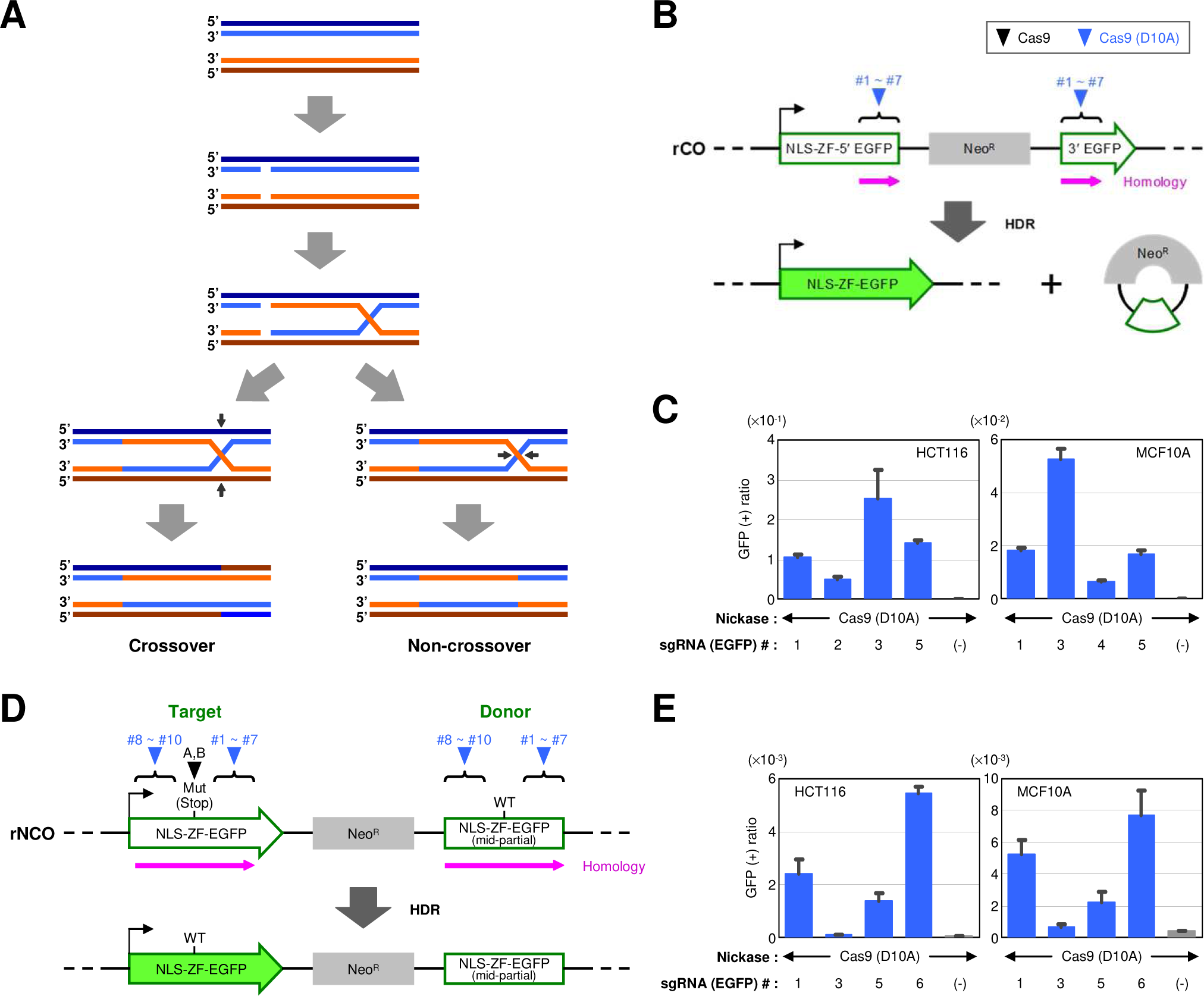
Tandem paired nicking leads to DNA recombination analogous to Holliday’s model. (A) Schematic description of Holliday’s model (Holliday, 1964; Liu and West, 2004). Lines depicted in four different colors represent DNA strands constituting two juxtaposed chromatids. Black arrows indicate cleavage in DNA strands that resolve Holliday junctions into crossover and non-crossover products. (B–E) Crossover and non-crossover recombination in the rCO (B and C) and rNCO (D and E) reporter systems, respectively. Cell clones harboring the rCO (B) or rNCO (D) reporter cassettes in the genome were transfected with a Cas9 nickase and analyzed by FCM. HDR between two EGFP fragments in the reporters results in restoration of a full-length wild-type NLS-ZF-EGFP gene. Purple arrows indicate the extent and direction of the homologous regions. NLS: nuclear localization signal, ZF: zinc finger, Mut (Stop): a mutation constituting an ectopic stop codon. Bar graphs indicate the ratios of GFP-positive cells representing crossover (C) and non-crossover (E) events. Mean and s.e.m. values of biological triplicates are shown. See also Figure S4.

We postulated that tandem paired nicking might also result in both crossover and non-crossover recombination. To investigate the outcomes of tandem paired nicking, we created cell clones that emit GFP signals upon crossover but not non-crossover recombination between two partial EGFP fragments (named the rCO clones; Figure 2B). When the same positions within two EGFP fragments were nicked by single Cas9 nickases, 0.7–25 % of the cells became GFP-positive (Figure 2C). To preclude the possibility that the restoration of a full-length EGFP was solely the outcome of a process similar to single-strand annealing elicited in these cells, we created an additional reporter clone in which one of the partial EGFP fragments was incorporated in the reverse orientation (named the rCOrev clone; Figure S4A). In this clone, a full-length EGFP gene should theoretically be restored by crossover but not single-strand annealing between EGFP fragments. Upon transfection of this clone with single Cas9 nickases against EGFP, we detected a modest increase in the percentage of GFP-positive cells approximately 2-log lower than those in the rCO clones (Figures S4B and S4C). Possible reasons for the large difference in the percentage of GFP-positive cells between the two reporter assays include distinct chromosomal locations of reporter integration in these clones, which may lead to differential accessibility of the reporters for Cas9 nickases. We subsequently performed flow cytometry (FCM)-based sorting of the GFP-positive cells from the rCOrev clone transfected with Cas9 nickases. An analytical PCR followed by sequencing of the PCR products obtained from the sorted cells verified the restoration of a full-length EGFP in these cells (Figure S4D).

We also employed cell clones that fluoresce when an EGFP reporter cassette in the genome undergoes non-crossover recombination with gene conversion but not crossover recombination (named the rNCO clones; Figure 2D). Transfection of this clone with single Cas9 nickases against EGFP resulted in the emergence of GFP-positive cell populations at up to 0.77 % (Figure 2E). Collectively, these data provide evidence that nicks introduced at the same position on two homologous DNA fragments lead to both crossover and non-crossover recombination.

### Donor length significantly affects the efficiency of targeted knock-in via tandem paired nicking

Given the genetic structure of the reporter cassette, the rNCO clones should be useful as platforms to assess the efficiency of targeted knock-in via tandem paired nicking. Indeed, transfection of this clone with two Cas9 nickases resulted in the emergence of GFP-positive cells, indicative of the correction of EGFP mutation via tandem paired nicking (Figures 3A and 3B). However, in this assay (hereafter called the rNCO assay), the efficiency of targeted knock-in via tandem paired nicking was significantly lower than that achieved via the Cas9 nuclease-based approach, in contrast to the results previously obtained for the *PIGA*-based assays. In an attempt to explain this difference, we performed the *PIGA* correction assay using a donor truncated from 1,991 bp to 603 bp in length and found that this truncation more profoundly affects the efficiency of targeted knock-in via tandem paired nicking than that achieved using a Cas9 nuclease (Figures 3C and 3D). These data suggest that the low efficiency of targeted knock-in via tandem paired nicking in the rNCO assay is at least partially due to the small size of homologous region (795 bp) in the reporter system. Other factors, such as distinct localization of donor DNAs between the rNCO assay and *PIGA*-based assays (chromosomal versus plasmid), might also affect the efficiency of targeted knock-in via tandem paired nicking.

**Figure 3.**
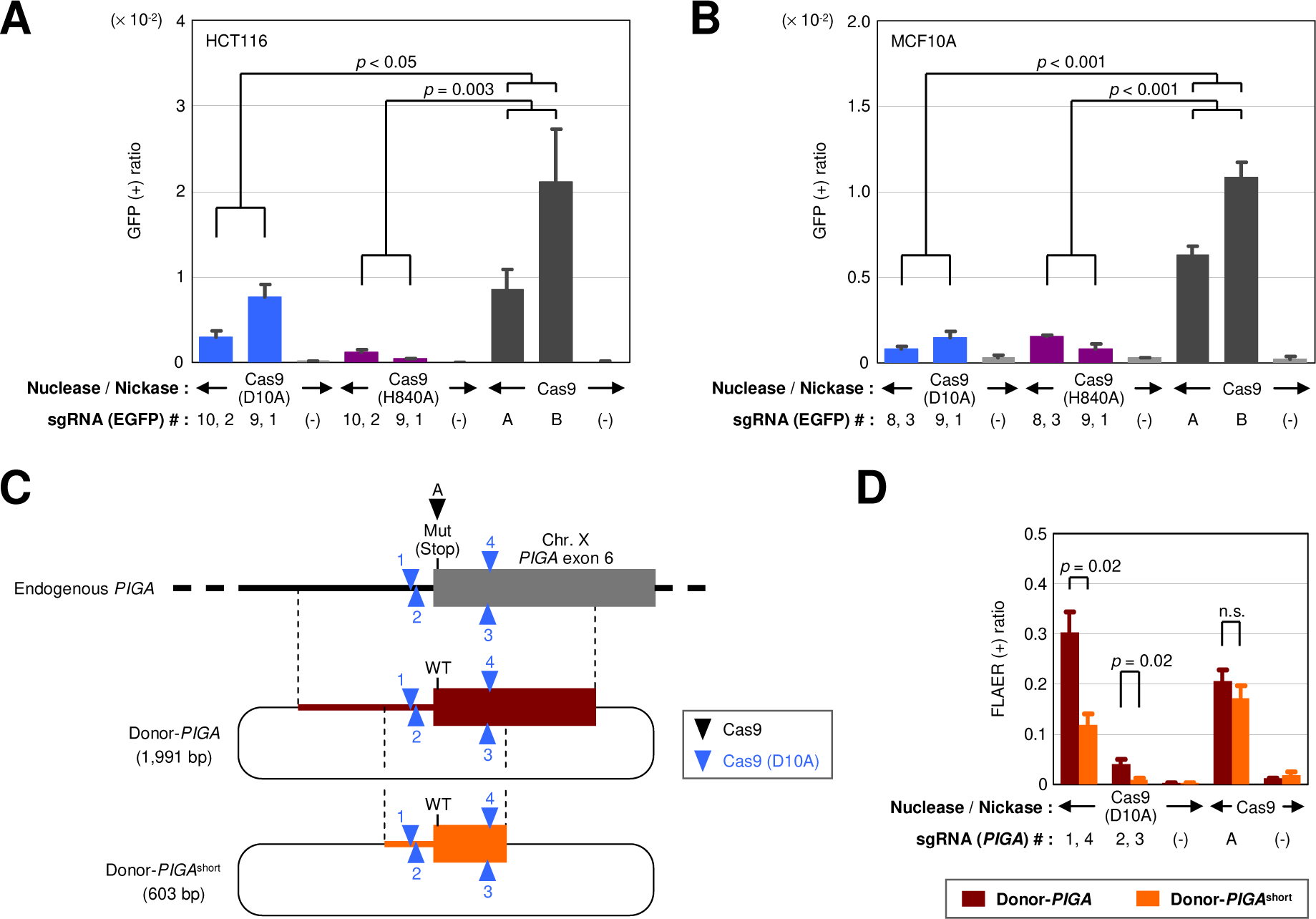
Donor length significantly affects the efficiency of targeted knock-in via tandem paired nicking. (A and B) Targeted knock-in in rNCO reporter clones (rNCO assay) mediated by tandem paired nicking. The rNCO clones generated from HCT116 (A) and MCF10A (B) were transfected with two Cas9 (D10A) nickases, two Cas9 (H840A) nickases, or a nuclease, and analyzed by FCM. Shown are the ratios of GFP-positive cells representing the occurrence of targeted knock-in (mean and s.e.m. of biological triplicates). (C and D) Scheme (C) and results (D) of the *PIGA* correction assay using a donor plasmid bearing a standard or a shortened homologous region. The length of homology is denoted below the name of the donor plasmid in (C). Mean and s.e.m. values of three independent experiments are shown in (D).

### Tandem paired nicking outperforms single nicking in targeted knock-in, particularly for targeted insertion of a large DNA fragment

Because transfection of single nickases into the rNCO clones led to the correction of an EGFP mutation via non-crossover type gene conversion (Figures 2D and 2E), targeted knock-in in endogenous genes might also be achievable using single Cas9 nickases together with a donor plasmid. We addressed this possibility by using *PIGA*-based assays and found that single nickases do mediate targeted knock-in (Figure 4). Overall, however, the use of single Cas9 nickases led to lower efficiency of targeted knock-in than tandem paired nicking, which was particularly evident in the RFLA performed with G418-based enrichment as shown in Figure 4B. In HCT116, the use of single Cas9 nickases resulted in virtually no targeted genomic integration of the Neo^R^ gene cassette (Figure 4B), while *PIGA* mutation was corrected with a decent efficiency using one out of four Cas9 nickases (Figure 4A). These data suggest that non-crossover-type gene conversion triggered by single Cas9 nickases may not be efficient enough to achieve targeted knock-in of large DNA cassettes, such as drug resistance gene cassettes, in many types of human cells. It is speculated that tandem paired nicking allows more efficient targeted genomic integration of a large DNA cassette by triggering two separate crossover events that encompass the DNA cassette.

**Figure 4.**
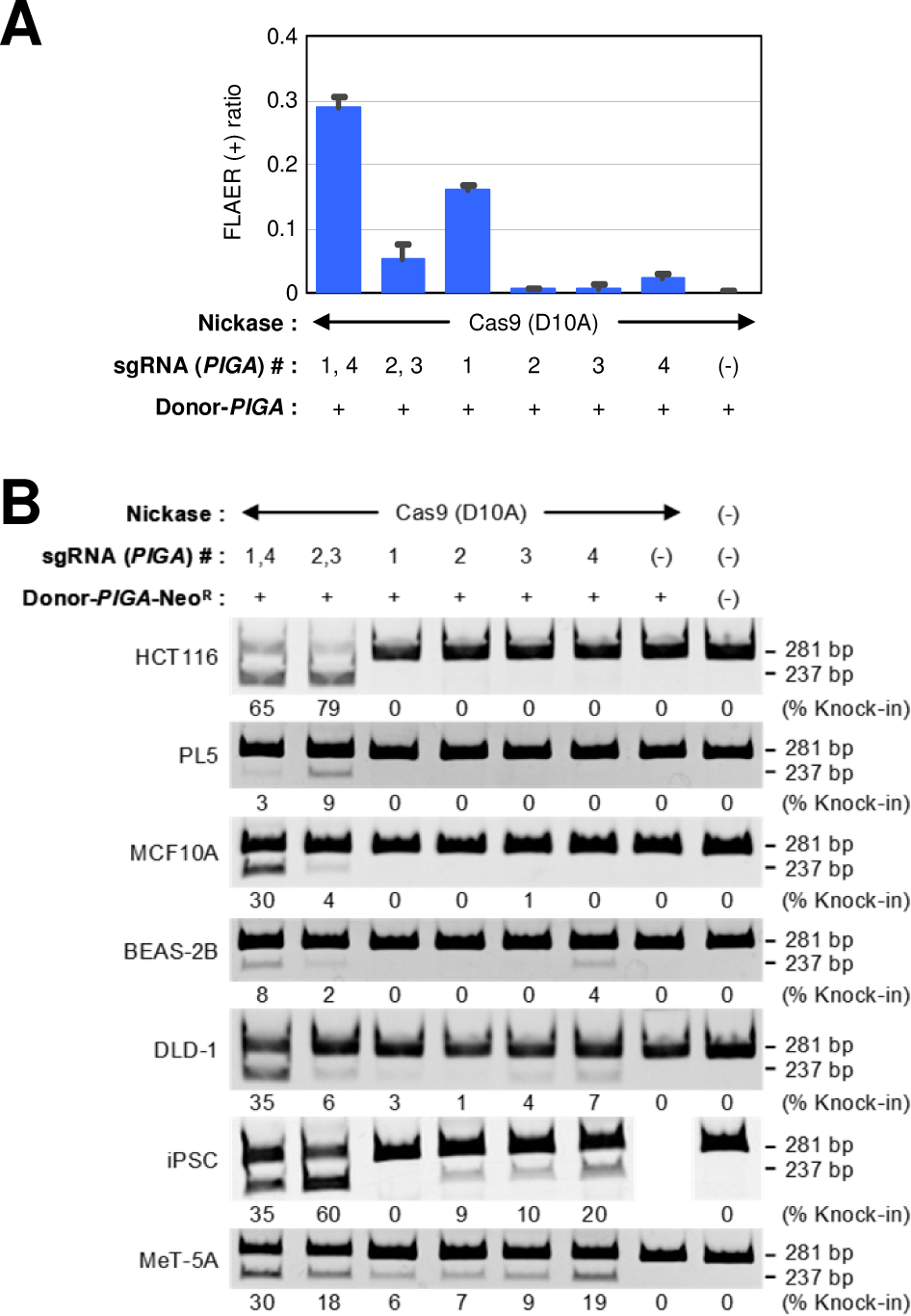
Tandem paired nicking outperforms single nicking in targeted knock-in, particularly for targeted insertion of a large DNA fragment. (A) *PIGA* correction assay using one or two Cas9 nickases along with Donor-*PIGA*. Mean and s.e.m. values of three independent experiments are shown. (B) Targeted knock-in of a *PIGA* mutation using one or two Cas9 nickases along with Donor-*PIGA*-Neo^R^. Percentages of knock-in alleles were calculated based on band intensities and are denoted under the corresponding lanes.

### Tandem paired nicking does not trigger a significant level of whole-donor targeted integration

If a crossover event is elicited by a single Cas9 nickase during targeted knock-in, a whole donor plasmid, including its backbone, should be integrated into the edited genome. In theory, such an event may occur regardless of whether targeted knock-in is conducted using one or two nickases. To address this issue, we conducted targeted *PIGA* disruption using one or two Cas9 nickases along with a donor plasmid, collected FLAER-negative cell populations by FCM-sorting, and performed Southern blot analysis (Figure 5A). This analysis provided no clear evidence for the targeted genomic integration of a whole donor plasmid that should be represented by a 10.8-kb band, suggesting that tandem paired nicking predominantly leads to legitimate ‘clean’ targeted knock-in (Figure 5B). A 4.3-kb band representative of an intact *PIGA* locus was observed for some of the samples that exhibited low knock-in efficiency. This band was present because FLAER-negative cells (*PIGA*-disrupted cells) are contaminated by a population of cells that bear no GPI-anchors, which sporadically emerges during cell culture due to spontaneous inactivation of *PIG* genes (including *PIGA*) that are essential for the biosynthesis of GPI-anchors.

**Figure 5.**
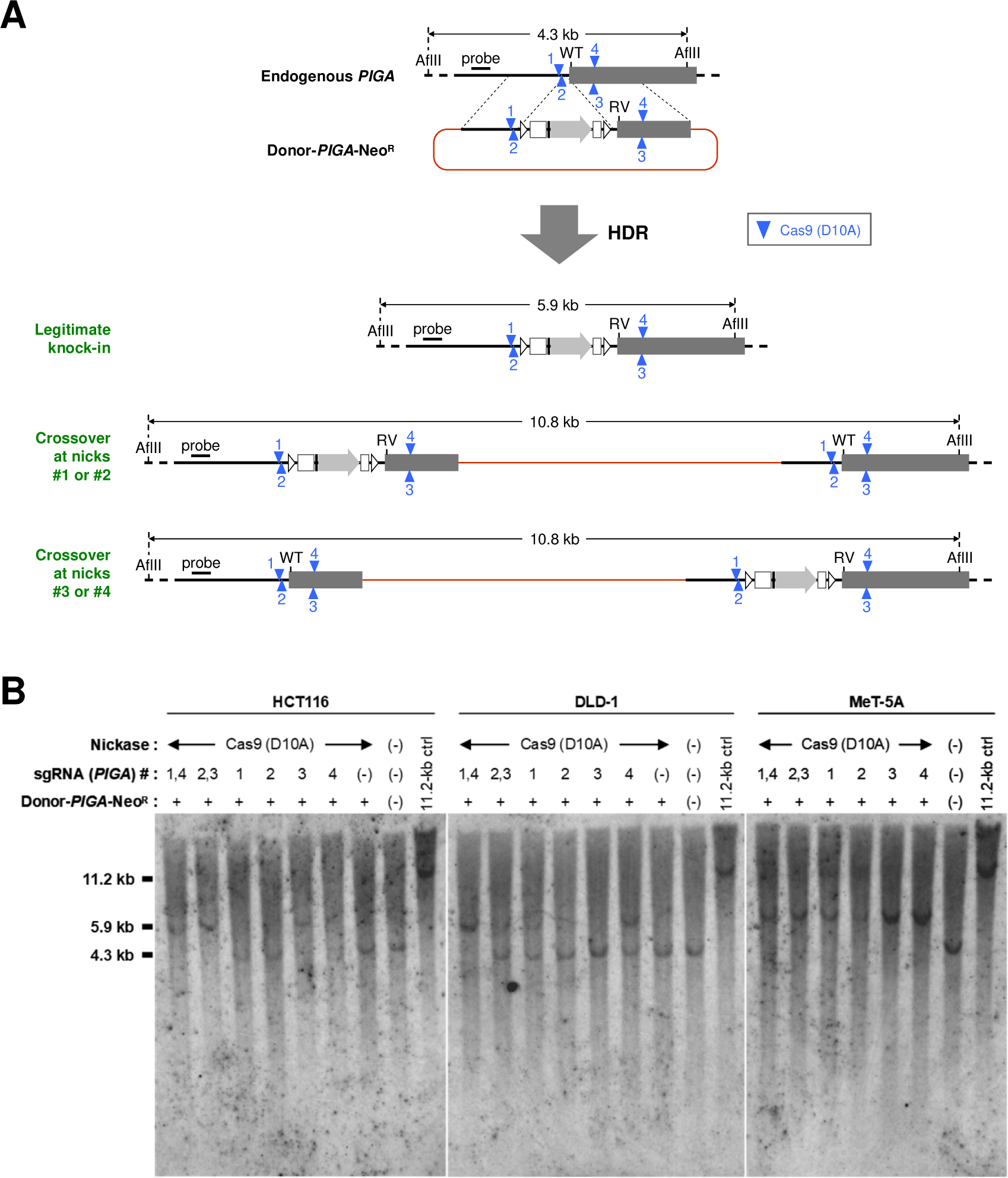
Tandem paired nicking does not trigger a significant level of whole-donor targeted integration. (A) Scheme for legitimate knock-in and whole-donor knock-in. For Southern blotting, parental cells were transfected with Donor-*PIGA*-Neo^R^ along with one or two Cas9 nickases, selected with G418, subjected to FCM-based sorting of FLAER-negative (*i.e.*, *PIGA*-disrupted) cells, propagated, and processed for Southern blot analyses using a probe shown in this scheme. AflII: recognition sites of AflII, a restriction enzyme used for Southern blot analysis. Brown lines represent the backbone of the donor plasmid. (B) Images of Southern blotting results. Approximate positions of the 11.2-kb, 5.9-kb, and 4.3-kb bands are marked on the left. gDNAs from untransfected parental cells were included in the analysis as controls. In addition, gDNAs from parental cells were double-digested with BamHI and SbfI and fractionated on agarose gels together with the other samples. Probing of these samples resulted in the formation of 11.2-kb bands, the appearance of which confirmed that the integrity of the gDNAs was sufficient for the formation of visible 10.8-kb bands. White lines within the gel indicate the borders of two distinct gels or two distinct portions of a gel. See also Figure S5.

We additionally performed *PIGA*-based assays using donor plasmids linearized by restriction enzyme digestion at sites on the plasmid backbone along with Cas9 nickases (Figure S5). Overall, the resulting data demonstrated that the efficiencies of targeted knock-in were similar to those obtained using uncut, circular donor plasmids (displayed in former figures) under corresponding conditions. This finding suggests the utility of both circular and linearized donor plasmids in targeted knock-in via tandem paired nicking.

### p53 does not significantly affect targeted knock-in via tandem paired nicking

It was recently shown that Cas9 nuclease-based genome editing triggers the activation of p53 signaling pathway and G1 cell cycle arrest. As targeted knock-in is mediated by HDR, which predominantly occurs in the S through G2 phases, Cas9 nuclease inherently obstructs targeted knock-in in a p53-dependent manner (Haapaniemi et al., 2018; Ihry et al., 2018).

Since tandem paired nicking elicits targeted knock-in without introducing appreciable DSBs, we reasoned that *p53*-dependent inhibition of targeted knock-in may be circumvented by the use of tandem paired nicking. Indeed, upon targeted knock-in of a *PIGA* mutation into MCF10A, tandem paired nicking did not significantly stimulate the expression of p21, a transcriptional target of p53, at both the messenger RNA (mRNA) and protein levels. In contrast, the use of Cas9 nucleases for transfection, instead of Cas9 nickases, significantly upregulated the expression of p21 mRNA and protein (Figures 6A–6D and S6). In addition, an rNCO assay with MCF10A revealed that the reversion of mutant EGFP to wild-type by tandem paired nicking was not significantly affected by the co-transfection of a short hairpin RNA against *p53*. The same *p53* suppression, however, significantly enhanced the EGFP reversion by Cas9 nucleases (*p* = 0.003; Figures 4E–4G). These data provide evidence that targeted knock-in via tandem paired nicking has little effect upon p53 activity, and is not significantly affected by wild-type p53, in contrast to the Cas9 nuclease-based approach. It is speculated, in general, that nick-based genome editing that create no DSBs during targeted knock-in may exhibit low level of reciprocal interference between p53 activity and targeted knock-in.

**Figure 6.**
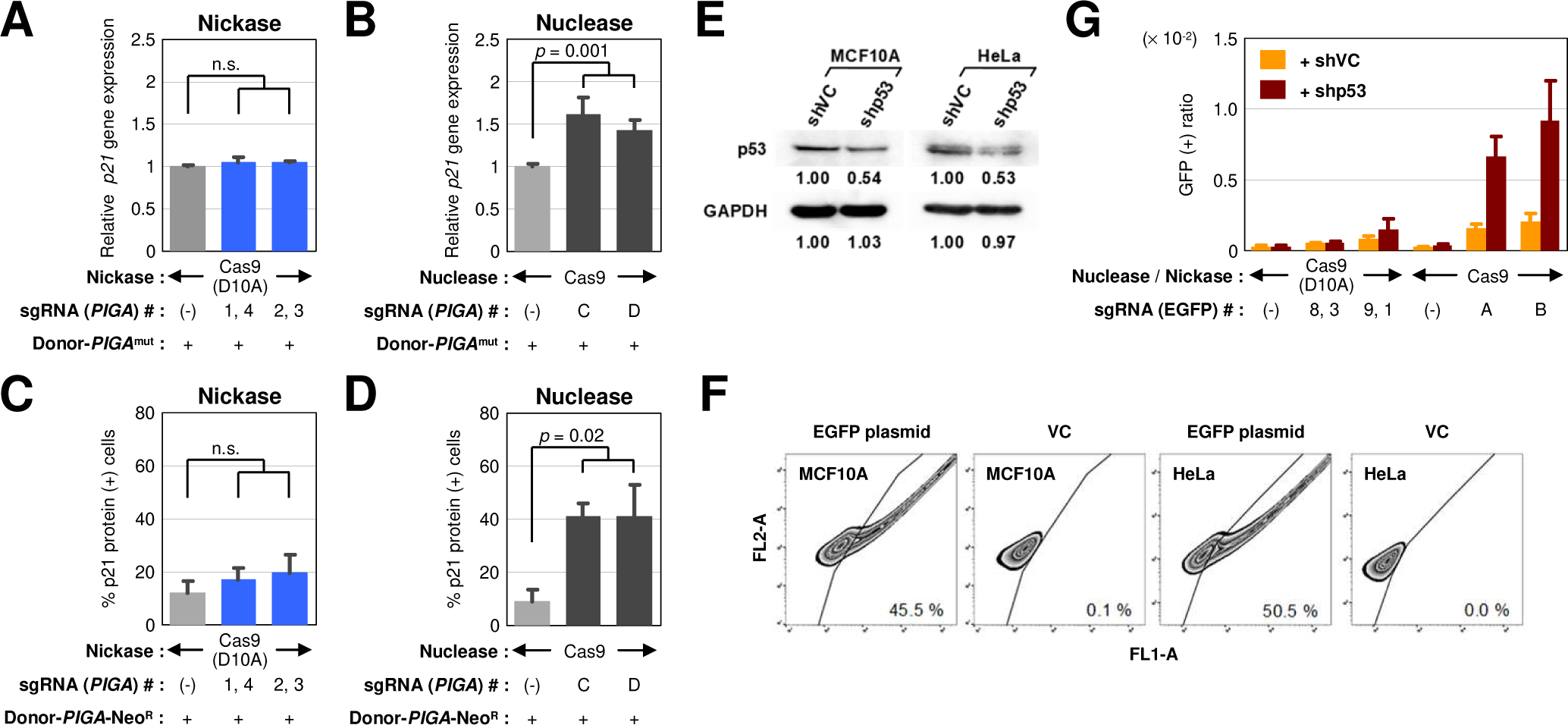
p53 does not significantly affect targeted knock-in via tandem paired nicking. (A and B) *p21* gene expression in the cells transfected with Cas9 nickases (A) or a nuclease (B). MCF10A cells were transfected with Cas9 nickases or a nuclease along with Donor-*PIGA*^mut^, and processed for real-time RT-PCR. The expression of *p21* is normalized to that of *GAPDH* and shown relative to the control samples (no sgRNA introduction). Mean and s.e.m. values of three independent experiments are shown. (C and D) Accumulation of the p21 protein in cells. MCF10A cells were transfected with Cas9 nickases (C) or a nuclease (D) along with Donor-*PIGA*-Neo^R^, immunostained with anti-p21 antibody and Alexa Fluor 594-labeled second antibody. EGFP expressed from the same plasmid with Cas9 and sgRNA was used as a Cas9 marker. Fluorescence signals emitted from the cells were quantified, and the ratios of p21-positive cells among GFP-positive cells were determined. G418 selection was not applied in this analysis. Mean and s.e.m. values of three independent experiments are shown. (E) Reduction in p53 levels by p53 shRNA. MCF10A and HeLa cells were transfected with a plasmid expressing an shRNA against p53 (shp53) or luciferase (shVC) and processed for Western blot analysis for detection of p53 and GAPDH. Band intensities relative to those in shVC transfectants are denoted under the lanes. (F) Transfection efficiencies in MCF10A and HeLa cells determined using a plasmid expressing EGFP. (G) rNCO assay in MCF10A. The MCF10A-rNCO clone was transfected with either Cas9 nickases or a nuclease, along with shp53 or shVC, and analyzed by FCM. Shown are the ratios of GFP-positive cells representing the occurrence of targeted correction of an EGFP mutation (mean and s.e.m. of three independent experiments). See also Figure S6.

### Targeted knock-in via tandem paired nicking is associated with infrequent indel formation at the edited genomic loci

The incidence of indel formation during targeted knock-in via tandem paired nicking has not been elucidated. To comprehensively understand the outcomes of targeted knock-in via tandem paired nicking, we introduced a *PIGA* ectopic stop codon into HCT116 and DLD-1 via either tandem paired nicking or Cas9 nuclease-based targeted knock-in (Figure 7A) and performed deep sequencing of a genomic region encompassing both the knock-in site and Cas9 target sites. Consistent with the results obtained in our previous *PIGA*-based assays, tandem paired nicking elicited targeted knock-in with efficiencies largely equivalent with those of Cas9 nucleases. Notably, only 2.4–4.2 % of the cells used for tandem paired nicking formed indels, in stark contrast with the Cas9 nuclease-based method, which generated indels in more than half of the cells (Figures 7B–7D). The ratios of targeted knock-in versus indel formation were 25–41-fold higher when tandem paired nicking rather than Cas9 nucleases was used (Figure 7E).

**Figure 7.**
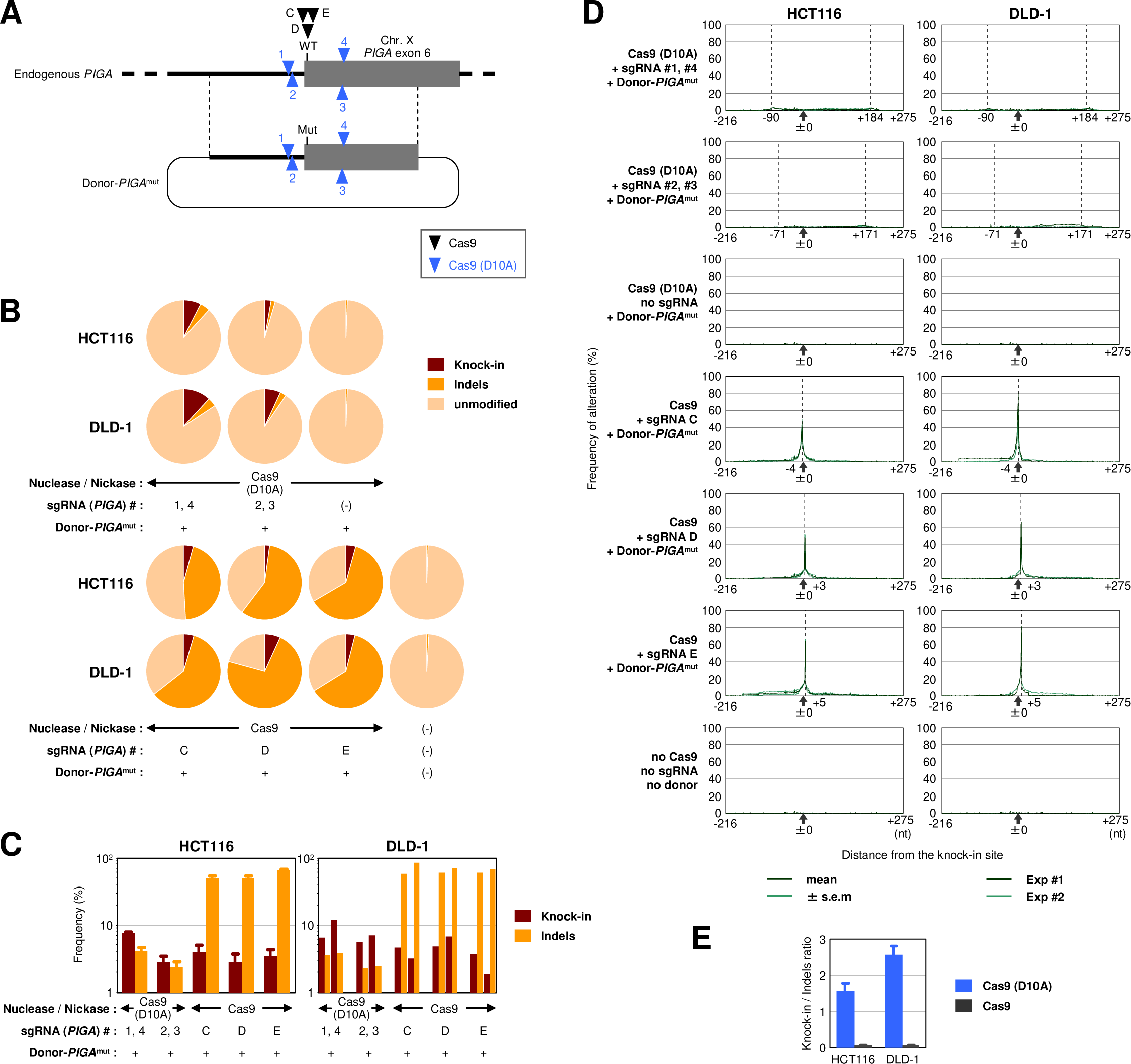
Targeted knock-in via tandem paired nicking is associated with infrequent indel formation at the edited genomic loci. (A) Scheme for the targeted knock-in of a *PIGA* mutation for which the incidence of indel formation was assessed. Cells were transfected with a Cas9 nuclease or nickases along with Donor-*PIGA*^mut^. The bulk populations of transfected cells were processed for PCR amplification of the *PIGA* targeted locus, followed by deep sequencing using a MiSeq sequencer. The data generated by MiSeq were analyzed using CRISPResso (Pinello et al., 2016). (B) Frequencies of targeted knock-in and indel formation in representative samples. (C) Graphical summary of the results. Mean and s.e.m. values of three independent experiments are shown for HCT116, while individual data obtained in two independent experiments are shown for DLD-1. Data for control samples were not included in (C) as these data were not appropriately presented in log scale. (D) Frequencies of nucleotide changes other than targeted knock-in at individual genomic positions surrounding the targeted site. Cas9 cleavage sites are indicated by vertical dotted lines. The values near the dotted lines with plus or minus signs represent the distances between the knock-in site (±0) and the Cas9 cleavage sites. (E) Ratios of targeted knock-in versus indel formation. Mean and s.e.m. values in Cas9 nickase- or nuclease-transfected samples are shown. See also Figure S7.

We also performed deep sequencing of the rNCO reporter cassette in HCT116 and MCF10A clones after eliciting HDR by tandem paired nicking or by using Cas9 nucleases. Consistent with previous data from rNCO assays (Figures 3A and 3B), the efficiency of targeted knock-in was higher when Cas9 nuclease rather than tandem paired nicking was utilized in this experiment. Meanwhile, the frequency of indel formation at the Cas9 target sites was 14–64-fold higher when Cas9 nucleases rather than tandem paired nicking were used (Figures S7A and S7B). As shown in Figure S7C, the ratio of targeted knock-in versus indel formation was 18-fold higher when tandem paired nicking rather than Cas9 nuclease was used (HCT116), or was similar between two approaches (MCF10A). Collectively, these data provide evidence that targeted knock-in based on tandem paired nicking is associated with less frequent indels around the targeted genomic loci than the Cas9 nuclease-based approach.

## Discussion

In this study, we demonstrated that concurrent nicking of a donor plasmid and the genome at the same position in the same strand (tandem paired nicking) enables targeted knock-in with a drastically reduced frequency of indels occurring at the edited genomic loci, yet with efficiency largely equivalent to that of the Cas9 nuclease-based approach. In addition, previous studies have repeatedly shown that indel formation at off-target genomic sites is markedly reduced by the use of Cas9 nickases rather than Cas9 nucleases (Cho et al., 2014; Ran et al., 2013a; Shen et al., 2014). These findings collectively provide evidence that tandem paired nicking permits targeted knock-in with improved fidelity and specificity as well as substantial efficiency equivalent to that of the previously described Cas9 nuclease-based approach.

We also demonstrated that wild-type p53 does not appreciably restrict targeted knock-in via tandem paired nicking. In contrast, when a Cas9 nuclease-based method was applied, targeted knock-in preferentially occurred in p53-suppressed cells as described previously (Haapaniemi et al., 2018; Ihry et al., 2018). A concern raised here is that, when using Cas9 nuclease, designed genetic modification might be preferentially introduced into cells with compromised p53 function, which can spontaneously emerge within cultured cells. If such preferential introduction occurs, the cells undergoing targeted knock-in may be more susceptible to transformation to cancer than their unmodified parental counterparts. However, this concern may not have direct clinical relevance, because gene-edited cells are subjected to careful examination to ensure safety prior to clinical use. Indeed, gene-edited cells have already been tested in clinical trials, and severe adverse effects have rarely been found (Cornu et al., 2017; Qasim et al., 2017; Tebas et al., 2014). Nonetheless, further safety in clinical gene editing will be established by developing a method of genome editing that hardly affects or is hardly affected by p53 activity. We propose that tandem paired nicking may help improve genome editing technology in this respect.

DNA recombination triggered by tandem paired nicking results in consequences analogous to those described in Holliday’s model (Holliday, 1964; Liu and West, 2004). It is now widely accepted that DSBs promote homologous recombination (HR) (Chapman et al., 2012; Wright et al., 2018). Meanwhile, our study demonstrated that iso-positional nicking of two homologous DNA fragments promotes a type of recombination that is similar to HR. Although Holliday’s model was proposed more than half a century ago, a direct assessment of whether this type of recombination truly occurs was recently made possible owing to the development of genome-editing tools, particularly, site-specific nicking enzymes. The molecular mechanisms underlying this recombination are largely unknown; however, another important consideration should be whether or not this recombination has physiological implications. It is currently unclear whether this recombination is a biological process that is inherent in cells or an incidental cellular response to artificial DNA lesions. One of the key steps for validation of this type of recombination as a physiological process will be the identification of intrinsic site-specific nickases in human cells. The identification of this type of recombination in other organisms is also warranted.

In a previous study, a relatively low efficiency of targeted knock-in via tandem nicking (tandem paired nicking) was documented (Nakajima et al., 2018). This modest efficiency of targeted knock-in via tandem nicking may be at least partially attributable to the length of homologous region. The reporter system used in this previous study had a short region of homology with a donor DNA consisting of an ATG-less EGFP gene (717 bp), which may have restrained targeted knock-in mediated by Cas9 nickases. We too employed an EGFP-based reporter system with a short homologous region (795 bp; rNCO assay) and observed low efficiencies of targeted knock-in via tandem paired nicking compared with those obtained by the Cas9 nuclease-based method. Several previous studies have described enhancement of targeted knock-in by creating two nicks, one on the donor and the other within the genome, at different positions (Chen et al., 2017; Davis and Maizels, 2014; Goncalves et al., 2012; Nakajima et al., 2018). This strategy, however, requires careful donor design, including the incorporation of a foreign sequence or silent mutation into the donor, which are eventually integrated into the genome upon targeted knock-in. In this regard, tandem paired nicking allows targeted knock-in to occur without additional nucleotide changes, including silent mutations. This property of tandem paired nicking enables precise and flexible genome engineering that introduces only desired genetic changes. Accordingly, for example, tandem paired nicking allows editing of gene promoters, enhancers, non-coding RNAs and intergenic regions. The editing of coding genes may also benefit from this precision, because additional nucleotide changes, including silent mutations, may affect transcription, splicing, translation and degradation of mRNAs derived from the target genes.

The distances between knock-in sites and Cas9 cleavage sites (hereafter called KC distances) in tandem paired nicking for *PIGA*, *CD55*, and the rNCO assay were 69–182 bp, 166–416 bp, and 116–217 bp, respectively, indicating that efficient targeted knock-in via tandem paired nicking can be accomplished with KC distances up to 416 bp. This result is in contrast with the Cas9 nuclease-based method, in which a KC distance larger than 20 to 30 bp markedly reduces the efficiency of targeted knock-in (Ran et al., 2013a). Thus, the use of tandem paired nicking offers better availability of Cas9 target sites within broader genomic windows than the Cas9 nuclease-based approach.

In summary, we have demonstrated that tandem paired nicking allows targeted knock-in with high precision, flexibility, and substantial efficiency. The cells subjected to genome editing via tandem paired nicking are likely to retain active p53 and, in the future, might offer low risk of carcinogenesis and high clinical utility. Further studies are warranted to better characterize this method and explore the applications of this method in various fields of life science.

## Materials & methods

### General molecular biological techniques

Genomic DNAs (gDNAs), except those used for Southern blotting, were extracted using the PureLink Genomic DNA Mini Kit (Thermo Fisher Scientific, Waltham, MA, USA). Extraction of total RNA and synthesis of complementary DNA (cDNA) were performed using the NucleoSpin RNA Kit (Takara Bio, Kusatsu, Japan) and High-Capacity cDNA Reverse Transcription Kit (Thermo Fisher Scientific), respectively. PCR was carried out using KAPA HiFi HotStart ReadyMix (Roche, Basel, Switzerland) and a Veriti thermal cycler (Thermo Fisher Scientific). For Sanger sequencing, the samples were prepared using the BigDye Terminator v3.1 Cycle Sequencing Kit (Thermo Fisher Scientific) and electrophoresed on a 3500 genetic analyzer (Thermo Fisher Scientific).

### Plasmids

The following plasmids were generously provided by researchers via Addgene: pSpCas9(BB)-2A-GFP (PX458) V2.0 (#48138), pSpCas9(BB)-2A-Puro (PX459) V2.0 (#62988), pSpCas9n(BB)-2A-GFP (PX461) V2.0 (#48140), and pSpCas9n(BB)-2A-Puro (PX462) V2.0 (#62987) from Dr. Feng Zhang (Broad Institute) (Ran et al., 2013b); pXCas9H840A (#60900) from Dr. Bruce R. Conklin (Gladstone Institutes) (Miyaoka et al., 2016); pCAG-EGxxFP (#50716) from Dr. Masahito Ikawa (University of Tokyo) (Mashiko et al., 2013); and pMKO.1 puro p53 shRNA 2 (#10672) from Dr. William Hahn (Dana-Farber Cancer Institute) (Masutomi et al., 2003).

The Cas9 (H840A) mutation (c.2635_2636CA>GC variation in the 3×FLAG-NLS-SpCas9 fusion gene) was transferred from pXCas9H840A to PX459, and the resultant plasmid expressing Cas9 (H840A) was named pH840Apuro. To construct sgRNA-expressing vectors, oligonucleotide pairs were designed (Table S1), annealed to each other, and cloned into PX458, PX459, PX461, PX462, or pH840Apuro as described previously (Ran et al., 2013b). The Donor-*PIGA*, Donor-*PIGA*-Neo^R^, Donor-*CD55*-Neo^R^, Donor-*PIGA*^short^, and Donor-*PIGA*^mut^ plasmids were generated by incorporation of the PCR-amplified *PIGA* or *CD55* gDNA fragments into relevant plasmid vectors followed by sequence verification of the amplified regions. Full sequences of the donor plasmids, along with information regarding the corresponding empty vector controls, are shown in Figure S8.

The reporter vectors used in the EGxxFP assays were constructed by PCR amplification of wild-type *PIGA* and *CD55* gDNA fragments using MCF10A gDNA as the template, followed by incorporation of these fragments into pCAG-EGxxFP.

The reporter vectors used to establish the rCO and rCOrev clones, namely, the prCO and prCOrev plasmids, respectively, were constructed by assembling genetic components originating from DR-GFP (Pierce et al., 1999) and PCR-amplified wild-type partial EGFP fragments within a retroviral backbone. The prNCO plasmid, used to produce the rNCO reporter clones, is identical to pBABE-HR, a generous gift from Dr. Ben H. Park (Vanderbilt University) (Konishi et al., 2011).

pMKO.1 puro p53 shRNA 2 was used to express a shRNA against p53 (shp53), while the corresponding control vector was constructed by removing the shp53 sequence from pMKO.1 puro p53 shRNA 2 and instead incorporating a shRNA against firefly luciferase (shVC). The sequences and features of key plasmids generated in this study are shown in Figure S8.

### Cell culture

All parental cell lines were obtained from American Type Culture Collection (ATCC; Manassas, VA, USA). HeLa and 293T cells were propagated in Dulbecco’s modified Eagle’s medium (D-MEM; Fujifilm, Tokyo, Japan) supplemented with 10 % fetal bovine serum (FBS; Merck, Darmstadt, Germany) and 1 % penicillin-streptomycin (P&S; Fujifilm). MCF10A cells and its derivative clones were maintained in D-MEM/Ham’s F-12 (Fujifilm) supplemented with 20 ng/ml epidermal growth factor (EGF; Merck), 500 ng/ml hydrocortisone (Merck), 10 µg/ml insulin (Fujifilm), 100 ng/ml cholera toxin (Merck), 5 % horse serum (Biowest, Nuaille, France), and 1 % P&S. HCT116 cells and its derivative clones were cultured in McCoy’s 5A (modified) medium (Thermo Fisher Scientific) supplemented with 5 % FBS and 1 % P&S. DLD-1, MeT-5A, and PL5 cells were propagated in Roswell Park Memorial Institute (RPMI) 1640 medium (Fujifilm) supplemented with 5 % FBS and 1 % P&S. BEAS-2B cells were maintained in Ham’s F-12 (Fujifilm) supplemented with 20 ng/ml EGF, 36 ng/ml hydrocortisone, 2.5 µg/ml insulin, 5 µg/ml transferrin (Merck), 130 pg/ml triiodo thyronine (T3) (Merck), 1 % FBS, and 1 % P&S. TIGE-9, a human induced pluripotent stem cell (iPSC) strain, was established previously (Shimojo et al., 2015), and maintained on mitomycin-C-treated SNL murine fibroblast feeder cells in D-MEM/Ham’s F-12 supplemented with 20 % knockout serum replacement (Thermo Fisher Scientific), 1 % non-essential amino acids (Merck), 0.1 mM 2-mercaptoethanol (Merck), 2 mM L-glutamine (Fujifilm), and 5 ng/mL basic fibroblast growth factor 2 (Peprotech, Rocky Hill, NJ, USA). The institutional ethical review committee at Aichi Medical University approved the protocol for the use of TIGE-9 cells in this study.

### Transfection

Transfection of plasmids was performed using the following devices and reagents according to the manufacturers’ instructions: 4D-Nucleofector System (Lonza, Basel, Switzerland) for HCT116, DLD-1, MeT-5A, PL5, and BEAS-2B cells; Neon Transfection System (Thermo Fisher Scientific) for TIGE-9 cells; Lipofectamine 3000 Reagent (Thermo Fisher Scientific) for MCF10A and HeLa cells; Polyethylenimine (PEI) Max (Polysciences, Warrington, PA, USA) for 293T cells. Derivative clones were transfected using the same reagents or devices as their parental cells.

In all assays involving plasmid transfection, HCT116, DLD-1, MeT-5A, PL5, and BEAS-2B cells (1 × 10^6^ cells) were electroporated with plasmids (1 µg each) and then seeded into a well of a 6-well tissue culture plate. TIGE-9 cells (2 × 10^5^ cells) were electroporated with plasmids (1 µg each) and then seeded onto feeder cells prepared in a well of a 6-well tissue culture plate. MCF10A, HeLa, and 293T cells were seeded into 6-well tissue culture plates at a density of 2 × 10^5^ cells/well and transfected with plasmids (2 µg each per well) on the following day.

### Correction of the *PIGA*-inactivating mutation (*PIGA* correction assay)

To create the HCT116-mut*PIGA* clone, HCT116 cells were transfected with Cas9 nickases along with Donor-*PIGA*^mut^, stained with FLAER (Brodsky et al., 2000) (Cedarlane, Ontario, Canada) approximately two weeks after transfection, and subjected to FCM-based sorting of FLAER-negative cells. Single cell-derived clones were developed from the sorted FLAER-negative cells, and subjected to RFLA screening using EcoRV. A cell clone in which an EcoRV recognition site was introduced was selected, and FLAER negativity was confirmed by FCM analysis. A genomic region flanking the knock-in site in the selected clone was PCR-amplified and sequenced to confirm the incorporation of the designed *PIGA*-inactivating mutation into the genome.

For the *PIGA* correction assay, the HCT116-mut*PIGA* clone was detached with Accutase (Innovative Cell Technologies, San Diego, CA, USA), and electroporated with PX459-, PX462-, or pH840Apuro-based constructs along with Donor-*PIGA*, Donor-*PIGA*^short^, or linearized Donor-*PIGA*. After transfection, cells were allowed to grow for three days, detached with Accutase, and stained with 0.1 µg/mL FLAER in PBS containing 0.25 mM EDTA and 0.4 % FBS. The percentages of FLAER-positive cells were determined by FCM analysis using a FACSCanto II system (BD Biosciences, Franklin Lakes, NJ, USA). Throughout this study, FCM analyses and cell sorting were carried out based on the initial FSC-A/SSC-A gating followed by FL1-A (530 ± 15 nm)/FL2-A (585 ± 21 nm) dot plots under 488-nm laser excitation.

In parallel with the other plasmids, PX458, a plasmid expressing EGFP, was transfected into the HCT116-mut*PIGA* clone together with Donor-*PIGA* in each experiment. The PX458-transfected cells and parent cell controls were analyzed by FCM on the following day. The percentage of GFP-positive cells in these samples was defined as the ʻtransfection efficiency’, and the percentage of FLAER-positive cells in the other samples was divided by this value, thereby providing the ʻFLAER-positive ratio’ for each sample.

### Targeted knock-in of inactivating mutations into the *PIGA* and *CD55* genes followed by G418 selection and RFLA

Cells were transfected with PX459- or PX462-based constructs along with Donor-*PIGA*-Neo^R^ or Donor-*CD55*-Neo^R^, and subjected to G418 selection for no less than two weeks. In some experiments, Donor-*PIGA*-Neo^R^ was used after linearization by PvuI cleavage of the backbone. The concentrations of G418 applied to cell lines were as follows: 0.12 mg/mL for MCF10A, 0.4 mg/mL for HCT116, 1.0 mg/mL for DLD-1, 0.8 mg/mL for MeT-5A, 0.5 mg/mL for PL5, 0.2 mg/mL for BEAS-2B, and 0.03 mg/mL for TIGE-9. Bulk populations of G418-resistant cells were collected using PBS containing 0.05 % trypsin and 0.02 % EDTA (BioConcept, Tallinn, Estonia) and processed for gDNA extraction for RFLA.

For RFLA, one of the PCR primers was placed distal to the homologous regions so that amplification could occur from both targeted and intact alleles but not from the donor DNA, and the knock-in site was included in the amplified region. The PCR primers used in these assays are listed in Table S2. PCR products were purified using the Gel/PCR DNA Isolation System (Viogene, New Taipei, Taiwan) and analyzed with a NanoDrop One instrument (Thermo Fisher Scientific) for determination of DNA concentrations. Three micrograms of the purified PCR products were then digested using a designated restriction enzyme and separated by TBE polyacrylamide gel electrophoresis, except for two experiments where a linearized donor DNA was used, for which agarose gels were used for electrophoresis. Digital images of the gels were obtained using an AE-6932GXES Printgraph (ATTO, Tokyo, Japan), and band intensities were quantified using ImageJ 1.52a. The percentages of *PIGA*-targeted alleles were obtained by the following equation: % *PIGA*-targeted allele = 28100 × ‘237-bp band’/(237 × ‘281-bp band’ + 281 × ‘237-bp band’). In the case of *CD55* editing, the following equation was used: % *CD55*-targeted allele = 24100 × ‘171-bp band’/(171 × ‘241-bp band’ + 241 × ‘171-bp band’). In both equations, the terms in quotation marks represent the intensities of the indicated bands.

For the experiments in which Southern blotting was performed after targeted knock-in of a *PIGA* mutation, bulk populations of G418-resistant cells were collected using Accutase and stained with 0.1 µg/mL FLAER in PBS containing 0.25 mM EDTA and 0.4 % FBS. FLAER-negative cells were sorted by FCM using a FACSAria III system (BD Biosciences), propagated in 75-cm^2^ flasks and used for Southern blot analyses with a probe complementary to a region in *PIGA* intron 5. The oligonucleotide primers used to amplify the probe sequence are listed in Table S2.

### IDAA

IDAA was performed as described previously (Carrington et al., 2015; Lonowski et al., 2017; Yang et al., 2015). Briefly, 293T cells were transfected with PX459-, PX462- or pH840Apuro-based constructs designed for *PIGA* or *CD55*, and gDNAs were extracted after three days of incubation. DNA fragments encompassing *PIGA* and *CD55* Cas9 target sites were amplified and labeled by nested PCR. The second (nested) PCR was carried out using three primers (0.025 µM non-labeled forward primer with a 5ʹ tail, 0.25 µM non-labeled reverse primer, and 0.25 µM 6-FAM-labeled tail primer named FamFwd). After the second PCR, 0.5 µL of 10 nM PCR product and 0.3 µL of GeneScan 500 LIZ Size Standard (Thermo Fisher Scientific) were mixed into 10 µL of Hi-Di Formamide (Thermo Fisher Scientific). The mixture was heated at 98 °C for 5 min, kept on ice for 5 min, and analyzed using a 3500 Genetic Analyzer and GeneMapper software version 5.0 (Thermo Fisher Scientific). Information regarding the PCR primers is provided in Table S2.

### EGxxFP assay

DLD-1 cells were transfected with pCAG-EGxxFP-*PIGA* or pCAG-EGxxFP-*CD55* along with PX459-, PX462-, or pH840Apuro-based constructs (Mashiko et al., 2013). Three days after transfection, cells were stained with 10 µg/mL Hoechst 33342 (Thermo Fisher Scientific), and fluorescent signals derived from EGFP and Hoechst 33342 were photographed using a BZ-9000 fluorescence microscope (Keyence, Osaka, Japan).

### Co-culture of *PIGA*- or *CD55*-positive and *PIGA*- or *CD55*-negative cells

Cell lines were plated onto 6-well tissue culture plates at approximately 50–70 % confluency and transfected with PX459- or PX462-based constructs shown in Tables S3 and S4 on the following day. In this assay, TransIT-LT1 reagent (Mirus Bio, Madison, WI, USA) was used for plasmid transfection as per the manufacturer’s instructions. Cells were selected with puromycin (HCT116, 0.4 µg/mL; DLD-1, 3 µg/mL; MCF10A, 1 µg/mL; PL5, 2 µg/mL; BEAS-2B, 2 µg/mL; MeT-5A, 1 µg/mL) from 48 hours through 96 hours post transfection and further propagated for no less than 10 days. After detachment of cells using Accutase, aliquots of the bulk cell populations were stained with FLAER (for *PIGA*-edited cells) or FITC-conjugated anti-human CD55 (IA10) mouse monoclonal antibody (BD Biosciences; for *CD55*-edited cells) and analyzed by FCM to determine the percentages of FLAER- or CD55-negative cells. The remainder of the cells were further propagated, and serial FLAER or anti-CD55 staining followed by FCM analysis was performed four times, with 3-days intervals, over the course of 13 days.

### GFP-based HDR assays using the rCO, rCOrev and rNCO clones

To establish rCO, rCOrev and rNCO reporter clones, HCT116 and MCF10A cells were transfected with the prCO, prCOrev, and prNCO plasmids, respectively, and single cell-derived clones were isolated. For HDR assays, the reporter clones were transfected with PX459- or PX462-based constructs designed for the NLS-ZF-EGFP gene. In the experiment that assessed the effect of p53 suppression, shp53 or shVC was co-transfected into the rNCO clone together with PX459- or PX462-based constructs. Three days after transfection, cells were treated with Accutase and suspended in PBS containing 0.25 mM EDTA and 0.4 % FBS. FCM analyses of the cells were performed to determine the percentages of GFP-positive cells using a FACSCanto II instrument. A subset of GFP-positive cells generated upon transfection of the rCOrev clone with PX462-based constructs were sorted using a FACSAria III instrument and propagated. gDNAs extracted from the resultant cells were used for analytical PCR followed by sequencing to verify the restoration of a full-length EGFP gene.

In parallel with the other plasmids, PX458, a plasmid expressing EGFP, was transfected into the reporter clones in each experiment. The cells electroporated and lipofected with PX458 were analyzed by FCM on the following day and three days later, respectively, together with the corresponding parent cell controls. The percentage of GFP-positive cells in these samples was defined as the ʻtransfection efficiency’, and the percentage of GFP-positive cells in the other samples was divided by this value, thereby providing the ʻGFP-positive ratio’ for each sample.

For Southern blotting, gDNAs derived from the rCO and rNCO clones were digested with either EcoRI or NheI, restriction enzymes whose recognition sites were absent in the prCO and prNCO plasmids, and processed for Southern blot analysis using a probe against the Neo^R^ gene. The oligonucleotide primers used to amplify the probe sequence are listed in Table S2.

### Southern blotting

Cells were propagated in 75-cm^2^ flasks at 80 % confluency prior to harvest. gDNA was extracted by standard phenol/chloroform extraction and ethanol precipitation. Eight micrograms of gDNA was digested with appropriate restriction enzymes, fractionated on 0.7 % agarose gels, and blotted onto a Hybond-N+ nylon membrane (GE Healthcare, Chicago, IL, USA). Probes were produced by PCR amplification and then labeled using the AlkPhos Direct Labelling Module (GE Healthcare). After hybridization, signals were visualized using CDP-Star Detection Reagent (GE Healthcare) and X-ray films (Fujifilm).

### Real-time reverse transcription-PCR (real-time RT-PCR)

MCF10A cells were transfected with PX459- or PX462-based constructs designed for the *PIGA* gene together with Donor-*PIGA*^mut^. Three days after transfection, total RNA was extracted from each transfectant and converted to cDNA. PCRs to amplify 296-bp and 185-bp DNA fragments from *p21* and *GAPDH*, respectively, were carried out in duplicate using SYBR Green I Nucleic Acid Gel Stain (Lonza) with a StepOnePlus instrument (Thermo Fisher Scientific). Three independent experiments were performed. For each PCR, a standard curve for each gene was generated using the data acquired with serially diluted samples, and gene expression levels were determined based on the standard curve. The expression levels of *p21* were graphically presented after normalization to those of *GAPDH*. Information regarding the primers used for amplification of *p21* and *GAPDH* is provided in Table S2.

### Immunostaining

MCF10A cells were grown on glass coverslips in 6-well tissue culture plates, transfected with PX458- or PX461-based constructs designed for the *PIGA* gene together with Donor-*PIGA*-Neo^R^, and incubated for three days. Although this donor plasmid harbors the Neo^R^ gene, G418 selection was not carried out in this assay. Cells were fixed with 4 % paraformaldehyde (Nacalai Tesque, Kyoto, Japan), permeabilized using PBS containing 0.1 % Triton X-100 (Nacalai Tesque), and blocked with PBS containing 7 % FBS. The cells were then incubated with anti-p21Waf1/Cip1 (DCS60) mouse monoclonal antibody (Cell Signaling Technology; CST, Danvers, MA, USA) and anti-GFP rabbit polyclonal antibody (Medical & Biological Laboratories, Aichi, Japan), followed by fluorescence staining with anti-mouse IgG Alexa Fluor 594-conjugated secondary antibody (Abcam, Cambridge, UK), anti-rabbit IgG Alexa Fluor 488-conjugated secondary antibody (Thermo Fisher Scientific), and Hoechst 33342. Finally, the cells were fixed on a glass slide using ProLong Glass Antifade Mountant (Thermo Fisher Scientific), and fluorescence signals were examined and photographed using a BZ-9000 fluorescence microscope. The captured images were analyzed using ImageJ 1.52a, and cells were designated as positive or negative for respective signals based on signal intensities. Three independent experiments were performed, and at least 25 (63.4 on average) GFP-positive cells per sample were assessed for p21 positivity.

### Western blotting

Plasmids expressing shRNA against p53 (pMKO.1 puro p53 shRNA 2; shp53) or luciferase (shVC) were transfected into MCF10A and HeLa cells. Three days later, cell lysates were harvested with Laemmli sample buffer, and protein concentrations were determined using the RC-DC Protein Assay (Bio-Rad Laboratories, Hercules, CA, USA). Equal quantities of protein were separated on SDS-polyacrylamide gels and transferred to PVDF membranes (Merck). The membranes were blocked with 1 % nonfat skim milk and incubated with anti-p53 rabbit polyclonal antibody (CST), anti-GAPDH (14C10) rabbit monoclonal antibody (CST), and anti-rabbit IgG HRP-linked antibody (CST). The reactivity was visualized by ImmunoStar LD (Fujifilm), and signals were captured and imaged using a Light-Capture II system (ATTO). Band intensities were quantified using ImageJ 1.52a.

To estimate the efficiency of shp53 transfection, MCF10A and HeLa cells were transfected with the pEGFP-IREShygro2 plasmid, and three days later, the percentage of GFP-positive cells was determined by FCM analysis. pEGFP-IREShygro2 (6,565 bp) was used as this plasmid was similar in length to the shp53 plasmid (6,370 bp).

### Deep sequencing of the targeted genomic loci

HCT116 and DLD-1 cells were transfected with PX459- or PX462-based constructs along with Donor-*PIGA*^mut^, while rNCO clones derived from HCT116 and MCF10A were transfected with PX459- or PX462-based constructs alone. After two weeks of incubation, gDNA was extracted, and nested PCRs were performed to amplify the targeted loci within *PIGA* or NLS-ZF-EGFP. For cell assays and gDNA extraction described above, two or three independent experiments for individual target loci and cell lines were performed. PCR and subsequent procedures, including library preparation for deep sequencing, were performed as a single experiment. For nested PCR, oligonucleotide primers were designed so that targeted genomic loci within *PIGA* and NLS-ZF-EGFP but not the donor DNA were amplified in the first round of PCR, and 491-bp and 545-bp fragments including knock-in sites and Cas9 target sites were amplified in the second round of PCR. For the second round of PCR, 33-bp and 34-bp adaptors were appended to the 5ʹ ends of the forward and reverse primers respectively, so that individual PCR products were tagged with unique indices during the third round of PCR (tagging PCR) using the Nextera XT Index Kit V2 (Illumina, San Diego, CA, USA). Information regarding the PCR primers is provided in Table S2.

After the second and third rounds of PCR, the PCR products were purified using Agencourt AMPure XP beads (Beckman Coulter, Indianapolis, IN, USA) according to the manufacturer’s protocol. After the third round of PCR, the quality and quantity of the PCR products were confirmed by agarose gel electrophoresis and measurement of spectral absorbance using a NanoDrop One instrument. Subsequently, the tagged PCR products (10 nM each) were pooled, and the solution was subjected to paired-end sequencing (351 + 251 cycles) using a MiSeq Personal Sequencer (Illumina) at the Advanced Genomics Center, National Institute of Genetics, Japan.

Paired-end reads produced by MiSeq were merged using PEAR (v0.9.6)(Zhang et al., 2014) with the default parameters. Alignment of merged reads with reference sequences as well as the calling of HDR and NHEJ reads was performed by CRISPResso (Pinello et al., 2016). Filtering of reads by average read quality (phred33 scale) greater than 30 and minimum single bp quality (phred33 scale) greater than 10 was applied for analysis with CRISPResso. The mean ± s.e.m. values of coverages were 56,000 ± 2,400, 58,000 ± 1,800, 3,100 ± 140, and 2,600 ± 220 for *PIGA* reads in HCT116, *PIGA* reads in DLD-1, rNCO reads in HCT116, and rNCO reads in MCF10A, respectively.

Among the four categories (‘HDR’, ‘Mixed HDR-NHEJ’, ‘NHEJ’ and ‘Unmodified’) to which aligned reads were sorted by CRISPResso, ‘HDR’ and ‘Mixed HDR-NHEJ’, where aligned reads carry the intended mutations introduced into the genome, were combined and are shown here as ‘knock-in’. The ‘NHEJ’ category from CRISPResso is shown here as ‘indels’. To obtain the frequencies of nucleotide changes other than targeted knock-in at individual genomic positions, the frequency of targeted knock-in was subtracted from the total mutation rates at the relevant genomic positions provided via CRISPResso.

### Statistical analyses

All statistical analyses were performed using Intercooled Stata (StataCorp, College Station, TX, USA). One-way repeated measures analysis of variance (ANOVA) with post hoc Bonferroni test was used to compare the efficiencies of HDR achieved by Cas9 (D10A), Cas9 (H840A), and Cas9 in the rNCO assay (Figures 3A and 3B) and to analyze the growth of co-cultured cells (Figure S3). Two-way repeated measures ANOVA was performed to compare the efficiencies of *PIGA* correction by Donor-*PIGA* and Donor-*PIGA*^short^ (two factors: ‘Cas9’ and ‘donor’; Figure 3D) and to analyze the effect of p53 suppression on the efficiency of targeted knock-in (two factors: ‘Cas9’ and ‘p53 status’; Figure 6G). As both analyses showed significant main effects for two factors as well as the existence of interaction, Bonferroni post hoc tests were performed at individual levels. The data for qRT-PCR and immunostaining-based assessment of p21 expression (Figures 6A–6D) were analyzed using the Wilcoxon rank-sum test. In all statistical analyses, a *p*-value less than 0.05 was considered to be significant.

### Contact for reagent and resource sharing

Plasmids for which sequences are provided in Figure S8 will be deposited to Addgene. Further information and requests for resources and reagents should be directed to and will be fulfilled by the Lead Contact, Hiroyuki Konishi.

### Data and software availability

All raw data from MiSeq sequencing have been deposited into the DNA Data Bank of Japan (DDBJ) under accession number DRA008492 (*PIGA*) and DRA008622 (rNCO).

## Acknowledgments

We would like to thank Mr. Makoto Naruse and colleagues at the Institute of Comprehensive Medical Research, Division of Advanced Research Promotion at Aichi Medical University, for providing technical assistance. We would like to thank Dr. Feng Zhang (Broad Institute) for providing the plasmids PX458, PX459, PX461, and PX462; Dr. Bruce R. Conklin (Gladstone Institutes) for pXCas9H840A; Dr. Masahito Ikawa (University of Tokyo) for pCAG-EGxxFP; Dr. William Hahn (Dana-Farber Cancer Institute) for pMKO.1 puro p53 shRNA 2; and Dr. Ben H. Park (Vanderbilt University) for pBABE-HR. This work was supported by Grants-in-Aid for Scientific Research (KAKENHI) from the Japan Society for the Promotion of Science (JSPS; 18K14703 to T.H., 18K08342 to A.O., 18H02645 to S.T., 17H05707 and 17K19465 to Y.O., 17K07263 to H.K.), Platform for Advanced Genome Science (PAGS; JSPS-KAKENHI 16H06279 to A.T.), Practical Research Project for Rare/Intractable Diseases from Japan Agency for Medical Research and Development (AMED; 19ek0109243h0003 to Y.O.), Hirose International Scholarship Foundation (to S.K.), and Takeda Science Foundation (to H.K.). M.L.R. is supported by the Japanese Government (MEXT) Scholarship for Research Students. M.W. was supported by the Nittoku Asia Foreign Student Scholarship.

## Author contributions

H.K. conceived, designed and supervised the project. H.K. designed the majority of the experiments, while T.H. designed the IDAA and EGxxFP assays as well as all the p53-related experiments. T.H. performed most of the experiments, while M.L.R. performed some of the GFP-based HDR assays, and S.K. performed initial experiments for the FLAER-based, RFLA-based and GFP-based assays. M.L.R. and S.K. created the new reporter clones. T.I. guided T.H. in the experiments using iPSCs. Y.O. supervised the iPSC experiments and offered all research equipment and reagents necessary to conduct these experiments. T.H. prepared samples for deep sequencing; A.T. performed MiSeq sequencing and initial processing of the data; and H.K. analyzed the resulting data with CRISPResso. T.H. and H.K. analyzed all the other data in the study. A.O., M.W., S.T., and Y.H. provided insights into the project. H.K. wrote the manuscript with significant contribution from T.H.

## Supplemental Information

**Figure S1.**
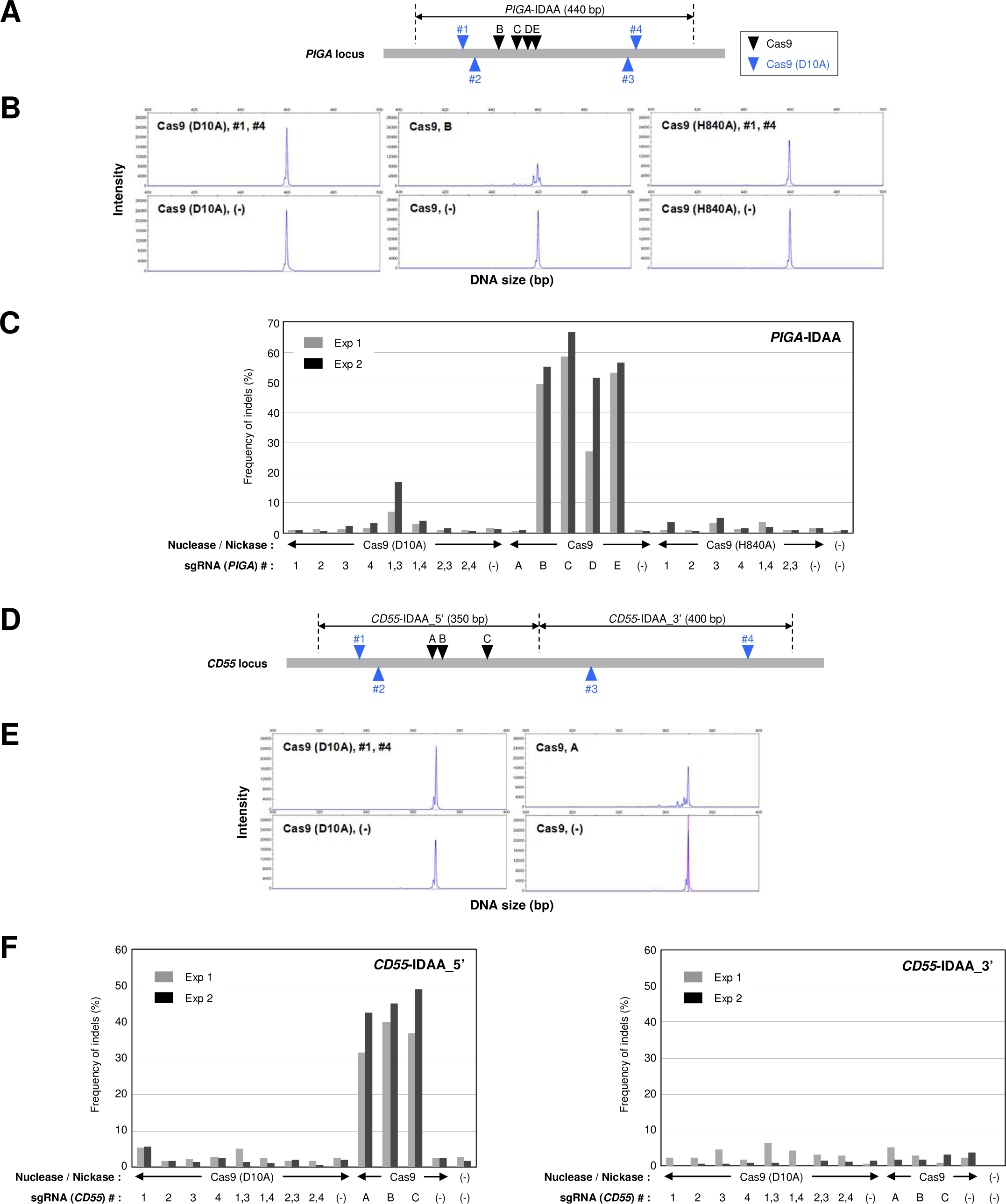
Tandem paired nicking does not elicit appreciable DSB. Related to Figure 1. 293T cells were transfected with Cas9 nucleases or nickases against *PIGA* (A–C) or *CD55* (D–F). The targeted genomic regions in *PIGA* (A) and *CD55* (D) were amplified by fluorescence PCR and subjected to capillary electrophoresis (IDAA assay). Representative chromatographs showing the intensities of the 400–500 bp (B) and 300– 400 bp (E) fluorescent PCR products, amplified from *PIGA* (B) and the 5ʹ side of *CD55* (E), respectively, are displayed. Bar graphs represent the summaries of IDAA results obtained in two independent experiments for *PIGA* (C), the 5ʹ side of *CD55* (F, left), and the 3ʹ side of *CD55* (F, right). Alphabetical and numerical identifiers in schematics (A) and (D) indicate the identical Cas9 nucleases and nickases as those illustrated in Figures 1B, 1E (*PIGA*), and 1I (*CD55*).

**Figure S2.**
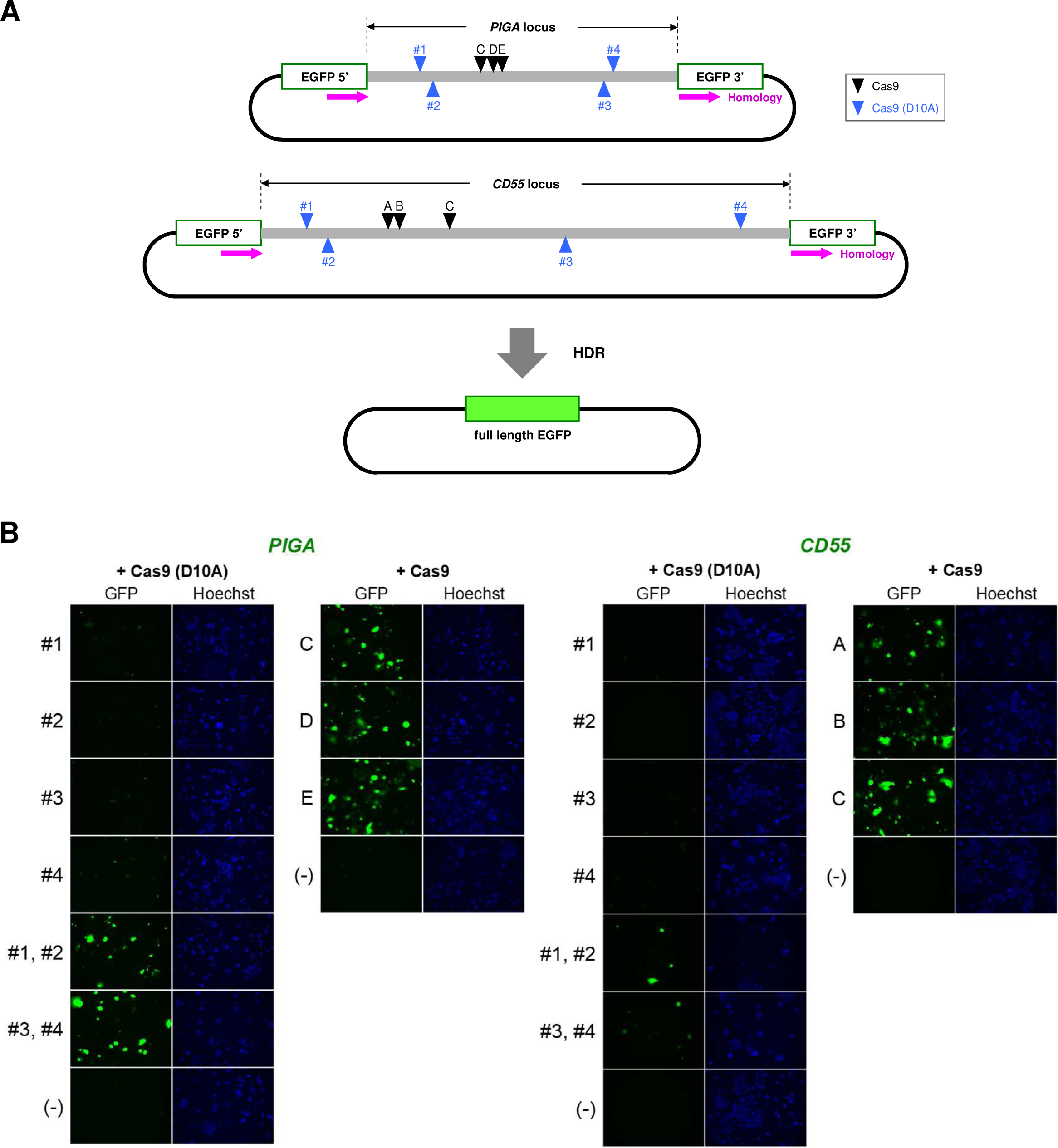
Cas9 nickases generate nicks in the *PIGA* and *CD55* sequences within transfected plasmids. Related to Figure 1. (A) DNA fragments derived from the corresponding gene loci were incorporated into pCAG-EGxxFP to create EGxxFP reporter plasmids. The resulting plasmids were transiently transfected into DLD-1 cells along with a Cas9 nuclease or nickases. DSBs in the plasmids lead to the reconstitution of a full-length EGFP via HDR (EGxxFP assay). Alphabetical and numerical identifiers in this schematic indicate the same Cas9 nucleases and nickases as those in Figures 1B, 1E (*PIGA*), and 1I (*CD55*). Purple arrows indicate the extent and direction of homologous regions. (B) Microscopic observation of the transfected cells. Representative images of GFP signals and nuclei are shown. Hoechst: Hoechst 33342.

**Figure S3.**
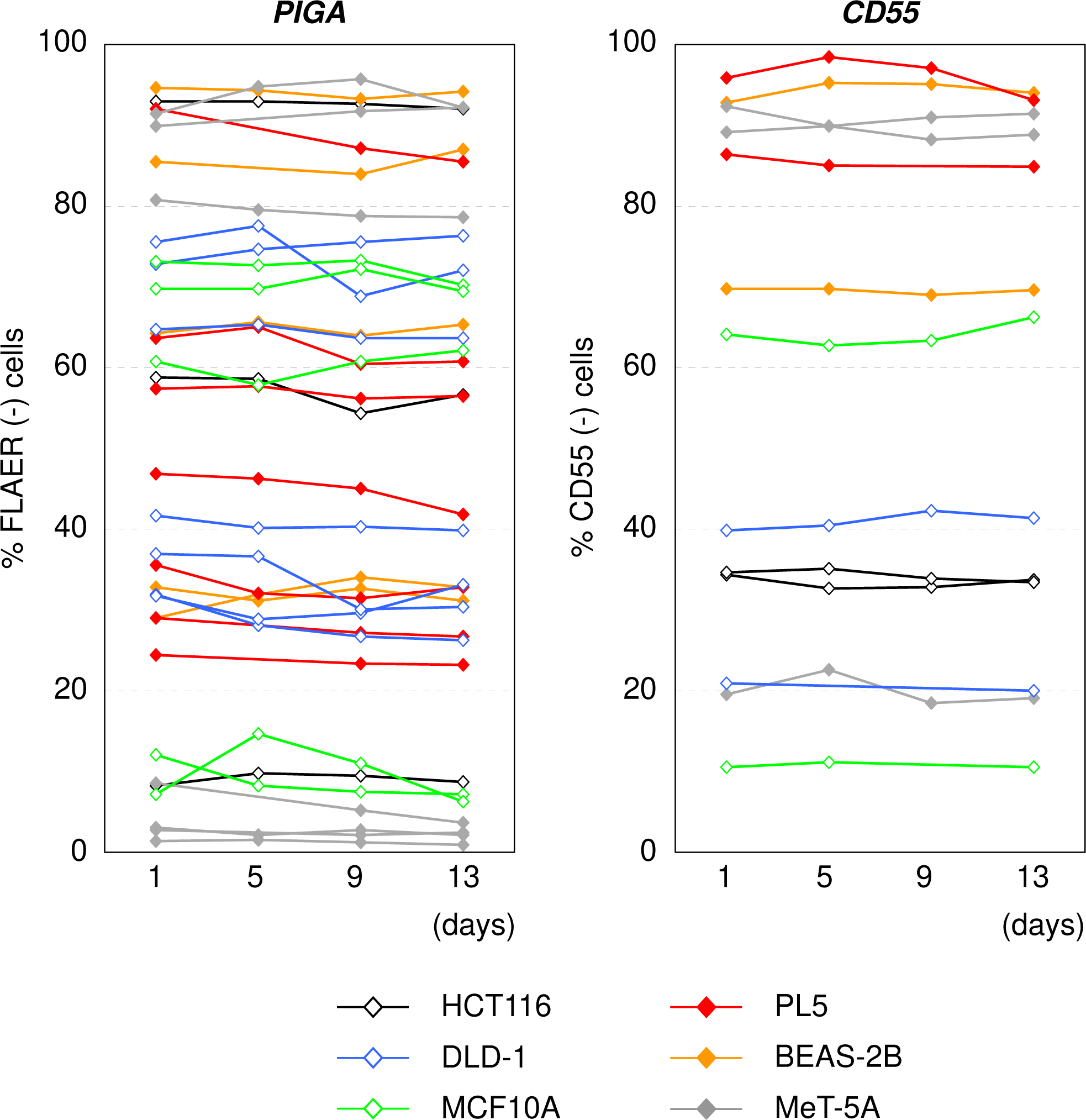
Time-course measurement of FLAER- and CD55-negative ratios in cultured cells exhibited no significant changes over time. Related to Figure 1. Cell lines indicated at the bottom were transfected with a Cas9 nuclease or paired Cas9 nickases against *PIGA* (left) or *CD55* (right). After at least two weeks of incubation, bulk populations of cells were stained with FLAER or fluorescence-conjugated anti-CD55 antibody and analyzed by FCM to determine the percentage of FLAER- or CD55-negative cells. For the same cell cultures, serial fluorostaining and FCM analysis of the cell aliquots were performed repeatedly over a 13-day course with 3-day intervals. Details regarding the Cas9 nucleases and nickases used for *PIGA* and *CD55* disruption are provided in Tables S3 and S4.

**Figure S4.**
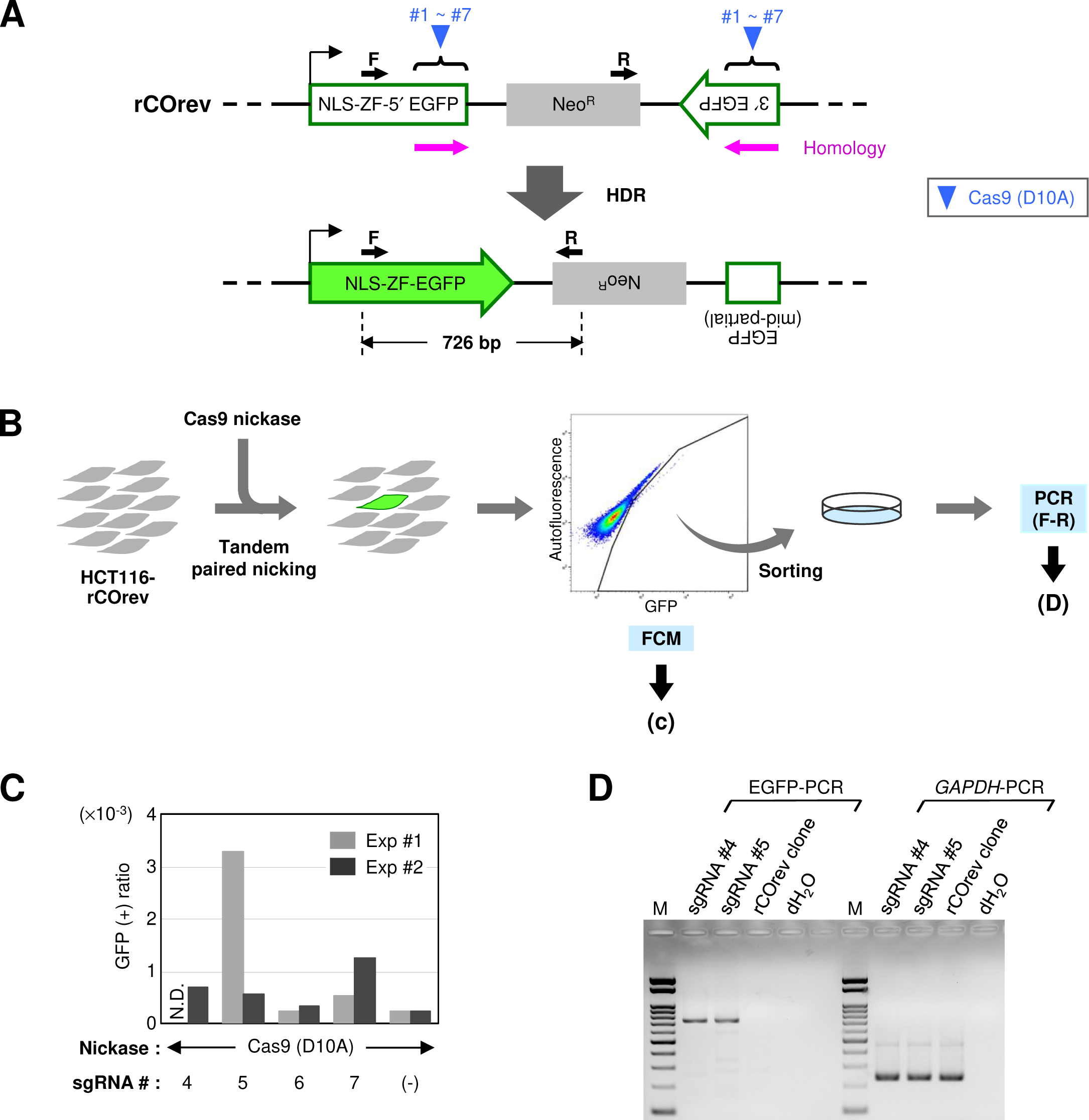
Tandem paired nicking triggers DNA crossover recombination. Related to Figure 2. (A) Scheme for the rCOrev reporter system. A cell clone harboring the rCOrev reporter cassette in the genome (top) was transfected with a Cas9 nickase against EGFP. A crossover event between two EGFP fragments in the reporter theoretically leads to the production of a full-length NLS-ZF-EGFP gene (bottom). Purple arrows indicate the extent and direction of the homologous region. (B) Experimental procedure. The rCOrev clone was transfected with Cas9 nickases and analyzed by FCM. GFP-positive cells were sorted, propagated, and processed for analytical PCR. (C) Ratios of GFP-positive cells representing the occurrence of HDR. N.D.: not done. (D) PCR products fractionated on an agarose gel. Portions of the EGFP and *GAPDH* genes were PCR-amplified using the sorted transfectants of indicated Cas9 nickases as templates. Primers depicted as F and R in (A) were used for PCR amplification of EGFP. gDNA from the rCOrev clone, in addition to distilled water, was used as a control for this analytical PCR. M: DNA size marker exhibiting 100–1,000-bp DNA fragments with an interval of 100 bp, plus 1,500-bp and 2,000-bp fragments.

**Figure S5.**
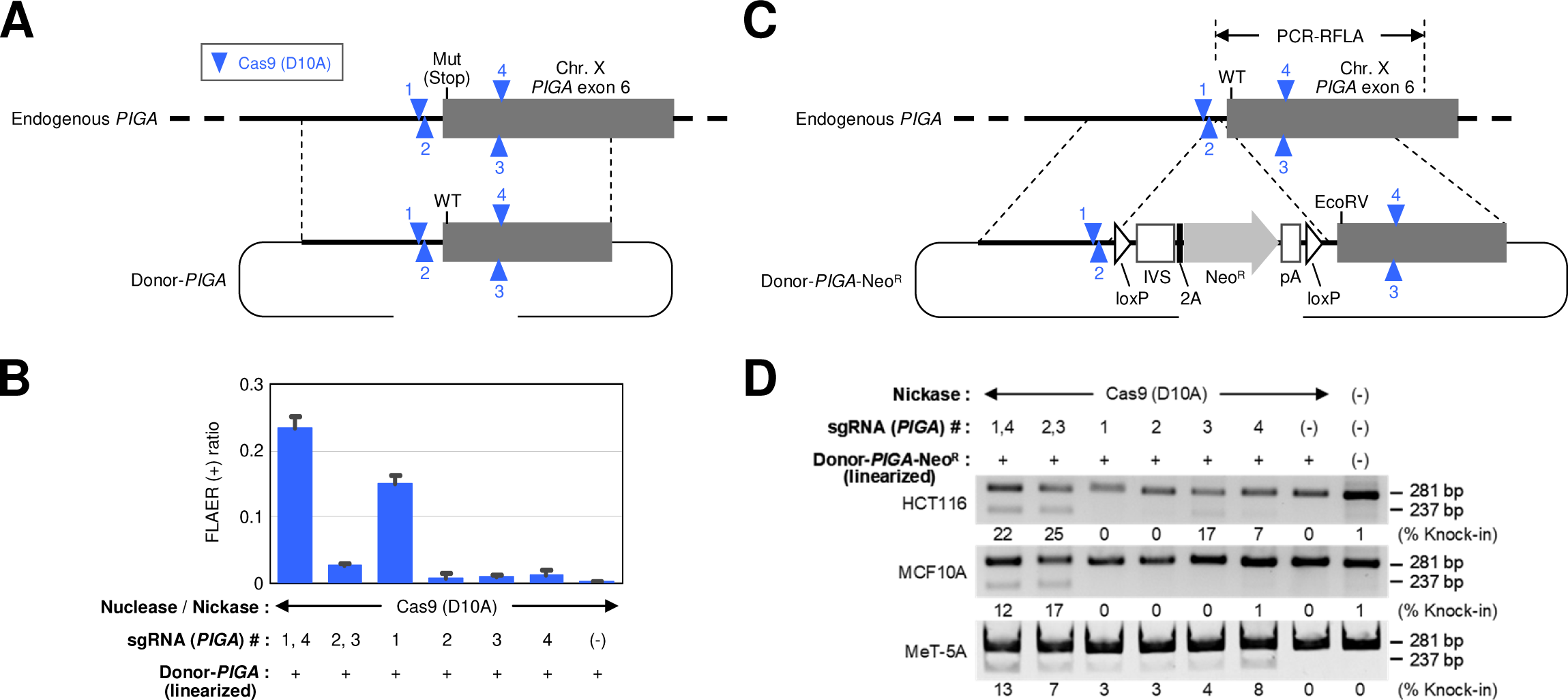
Linearized donor plasmids support targeted knock-in via tandem paired nicking. Related to Figure 5. (A) *PIGA* correction assay using a linearized donor plasmid. Donor-*PIGA* was linearized at the backbone by BsaI digestion and transfected into the HCT116-mut*PIGA* clone together with one or two Cas9 nickases. Cells were then stained with FLAER and analyzed by FCM. (B) Ratios of FLAER-positive cells representing the occurrence of *PIGA* correction. Mean and s.e.m. values of three independent experiments are shown. (C) Targeted knock-in of a *PIGA* mutation using a linearized donor plasmid. Donor-*PIGA*-Neo^R^ was linearized by cleavage with PvuI at the backbone and transfected into cell lines together with one or two Cas9 nickases. Cells were then selected with G418 and processed for RFLA. (D) Gel images for RFLA. Percentages of knock-in alleles were calculated based on band intensities and marked under the corresponding lanes.

**Figure S6.**
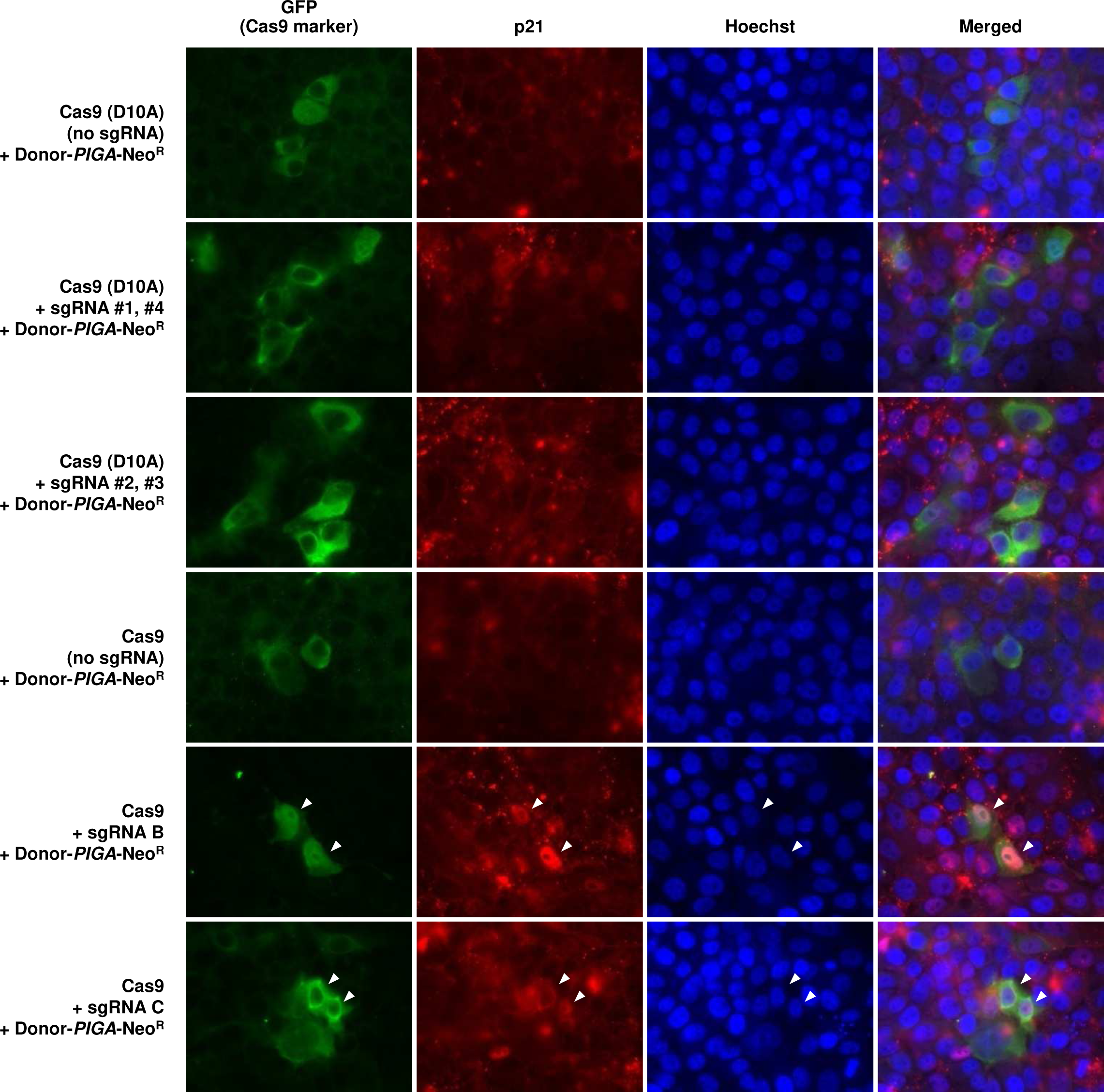
Representative microscopic images for p21 immunostaining. Related to Figure 6. GFP signal (enhanced by Alexa Fluor 488-labeling) indicates cells expressing Cas9. Cells expressing p21 were visualized by Alexa Fluor 594. White arrowheads indicate the cells that were designated as positive for both GFP and Alexa Fluor 594. Hoechst: Hoechst 33342. Merged: merged images of GFP, Alexa Fluor 594, and Hoechst 33342.

**Figure S7.**
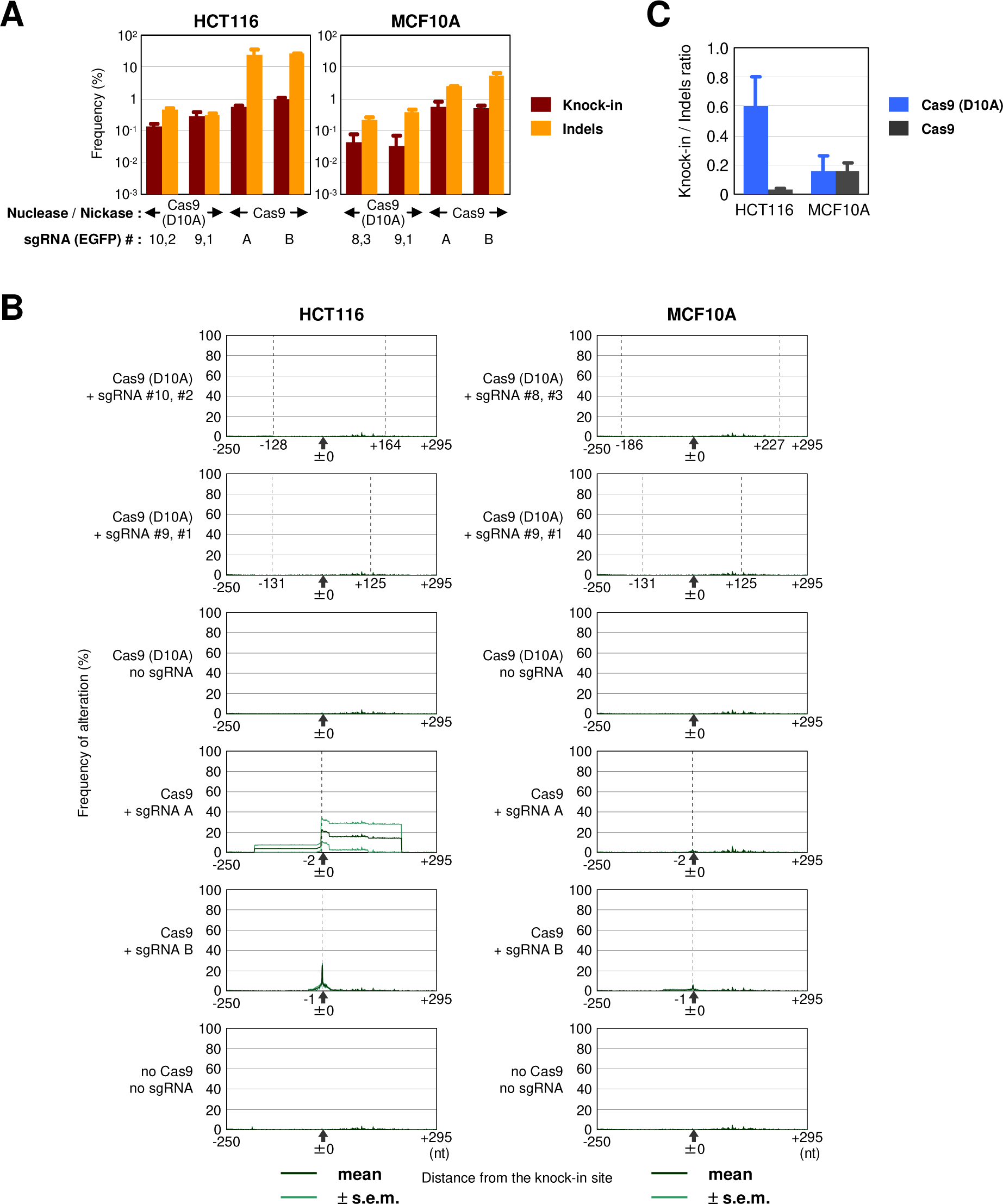
HDR within the rNCO reporter cassette via tandem paired nicking is associated with infrequent indel formation. Related to Figure 7. (A) Frequencies of HDR within the rNCO reporter cassette (targeted EGFP correction) and indel formation elicited by Cas9 transfection. rNCO clones derived from HCT116 and MCF10A were transfected with a Cas9 nuclease or nickases. The bulk populations of transfected cells were processed for PCR amplification of the rNCO reporter cassette, followed by deep sequencing using a MiSeq sequencer. Mean and s.e.m. values of three independent experiments are shown. Data for control samples are not included in (A) as these data are not appropriately presented in log scale. (B) Frequencies of nucleotide changes other than targeted EGFP correction at individual genomic positions surrounding the EGFP correction site. Cas9 cleavage sites are indicated by vertical dotted lines. The values near the dotted lines with plus or minus signs represent the distances between the correction site (±0) and the Cas9 cleavage sites. (C) Ratios of EGFP correction versus indel formation. Mean and s.e.m. values for Cas9 nickase- and nuclease-transfected samples are shown.

**Figure S8.**
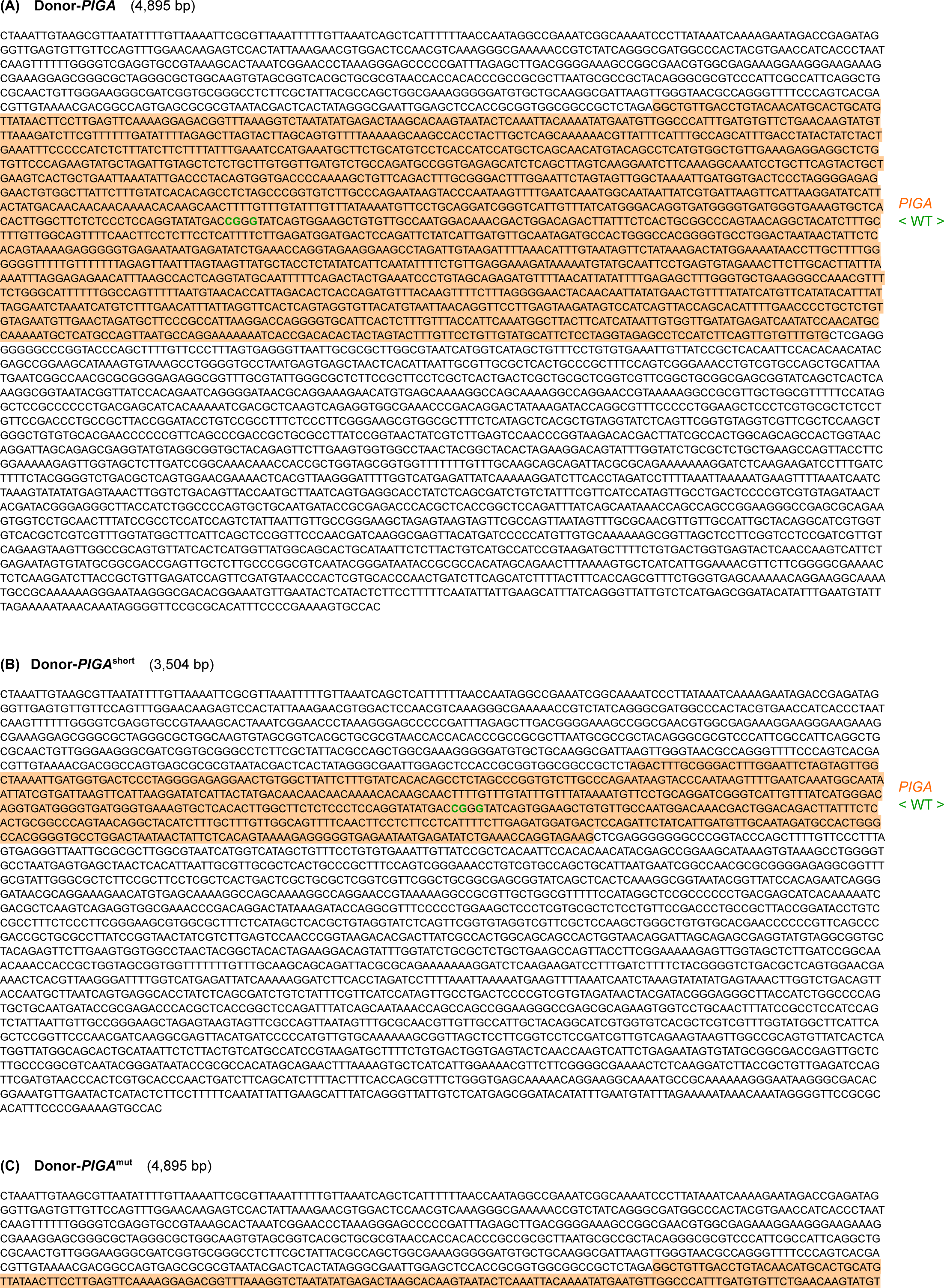
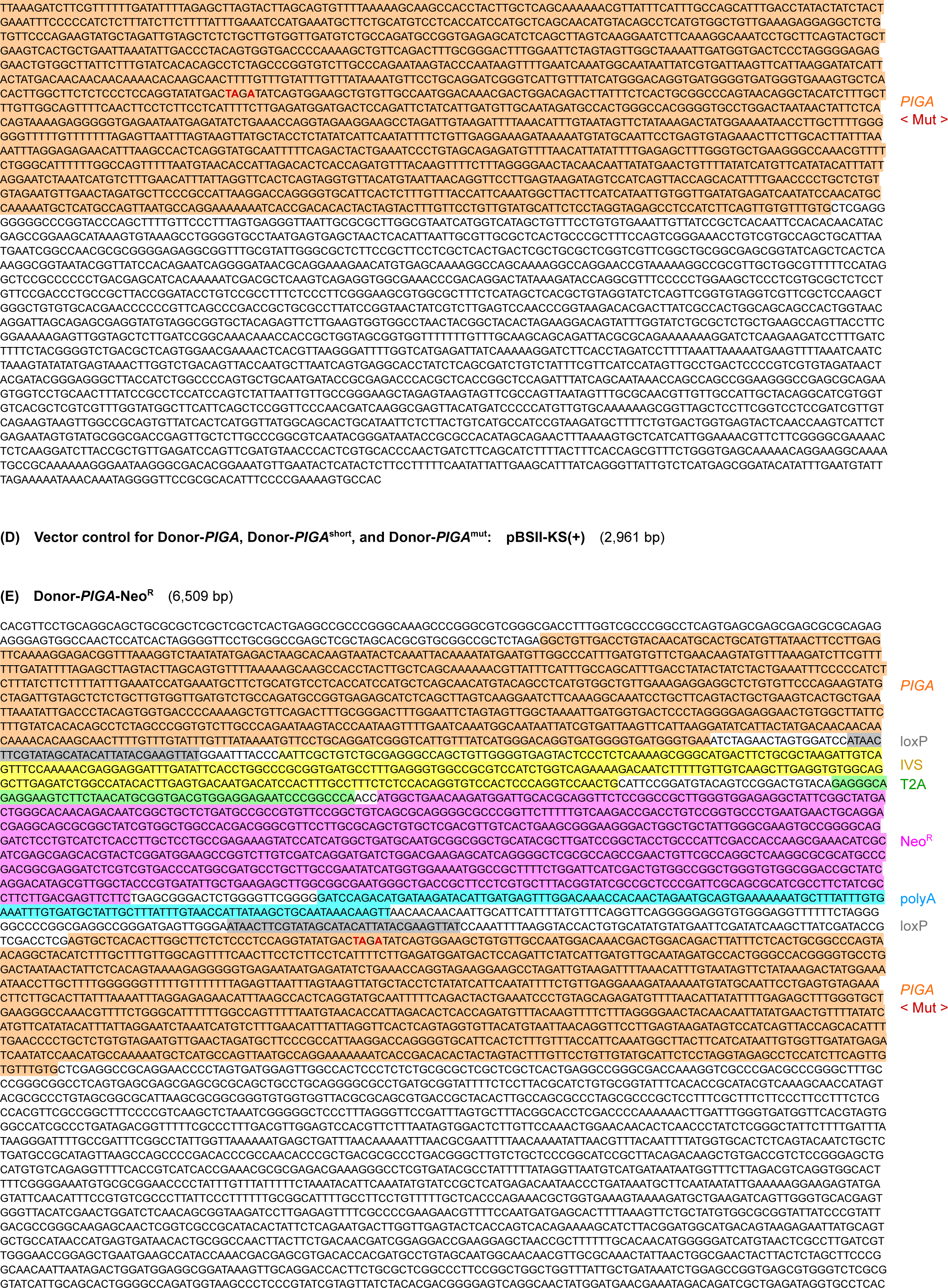
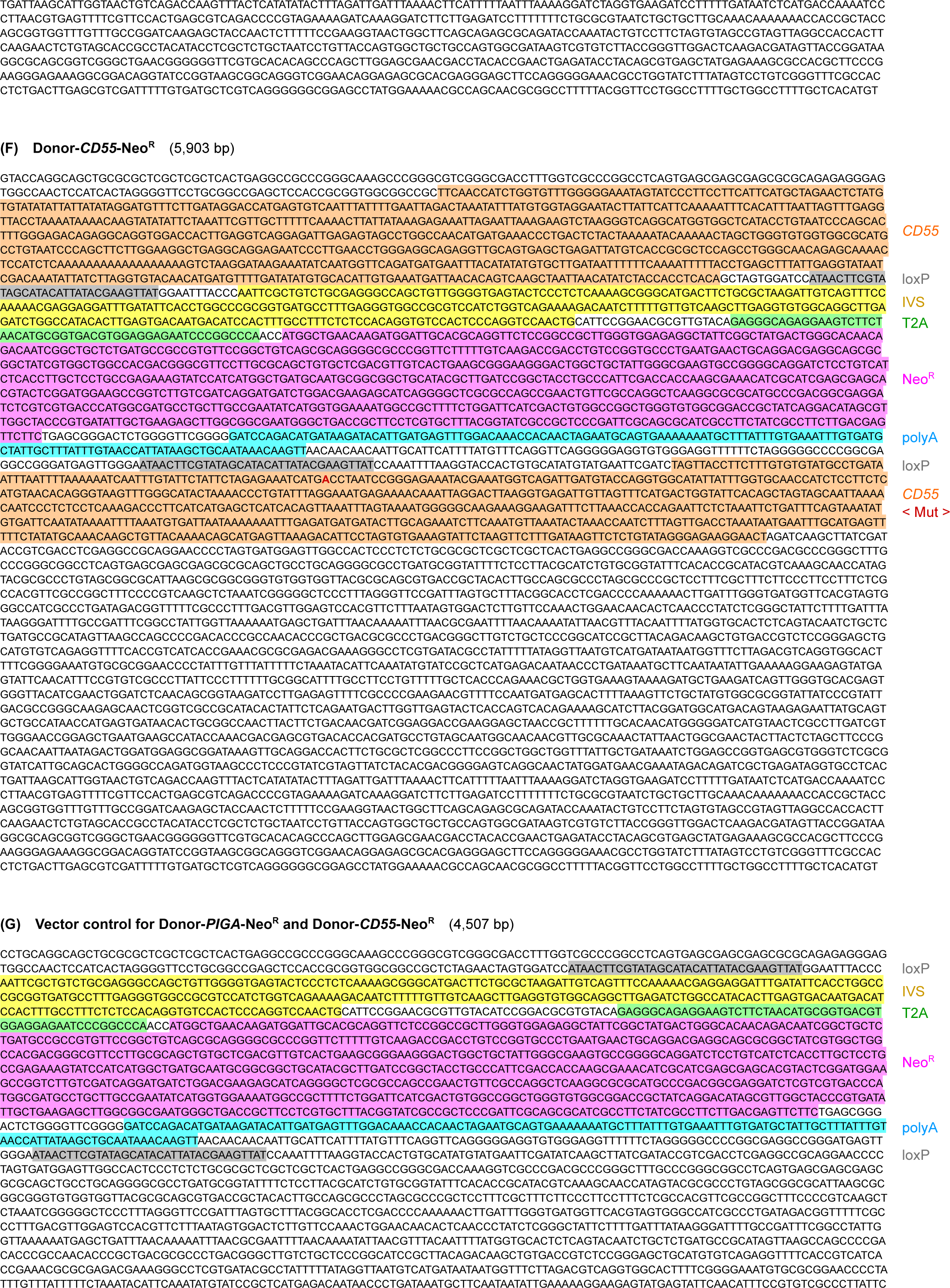
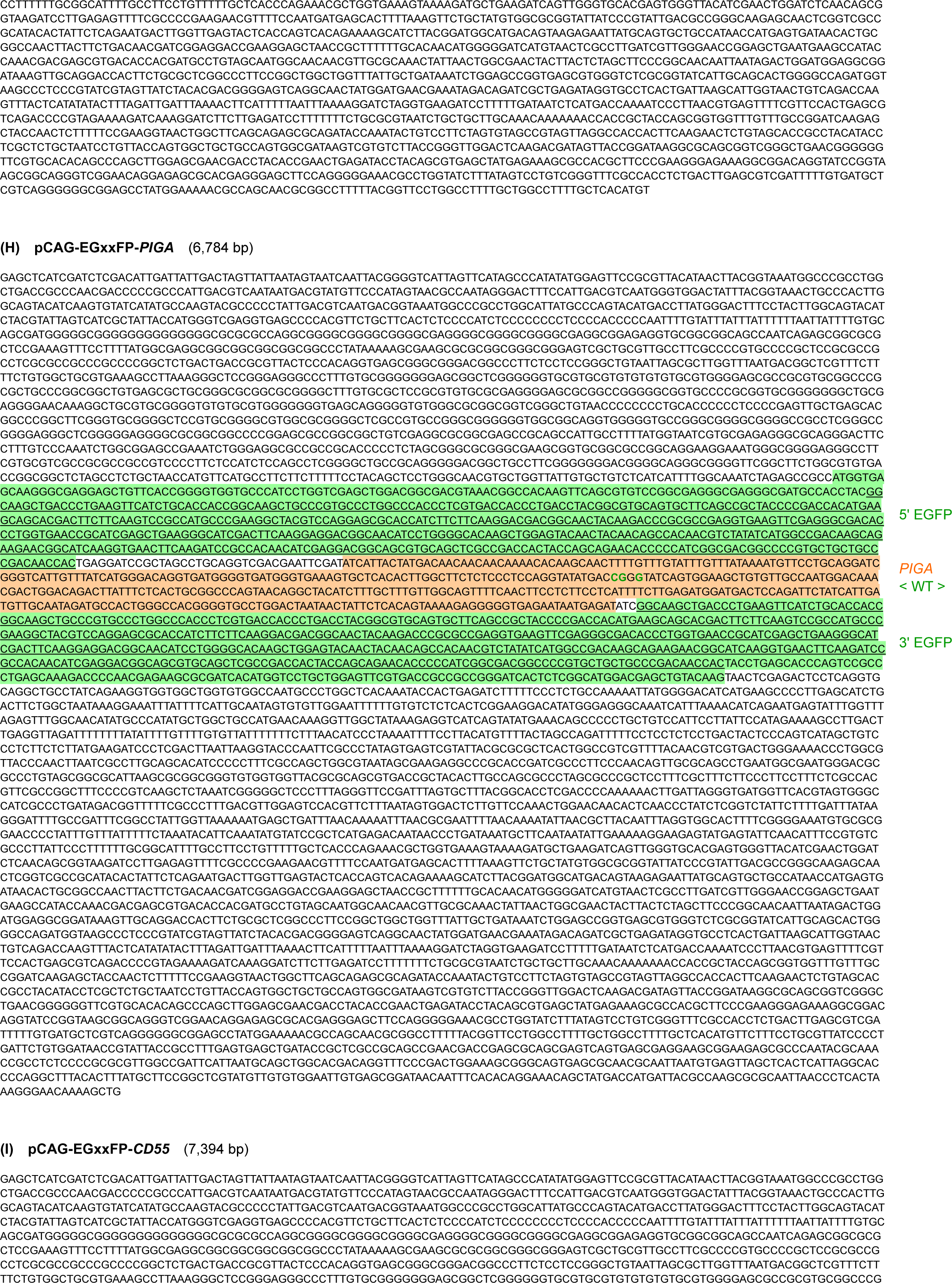
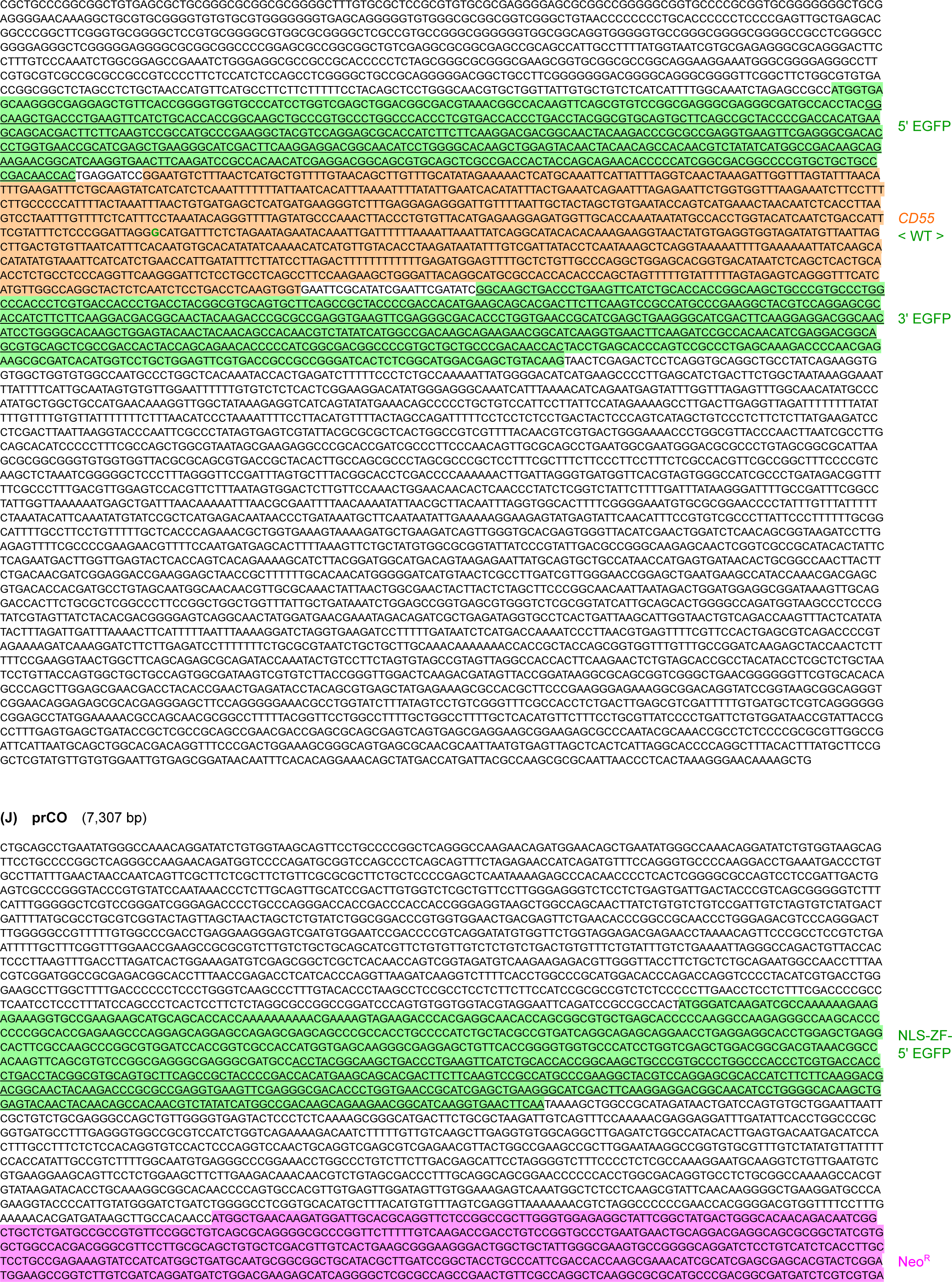
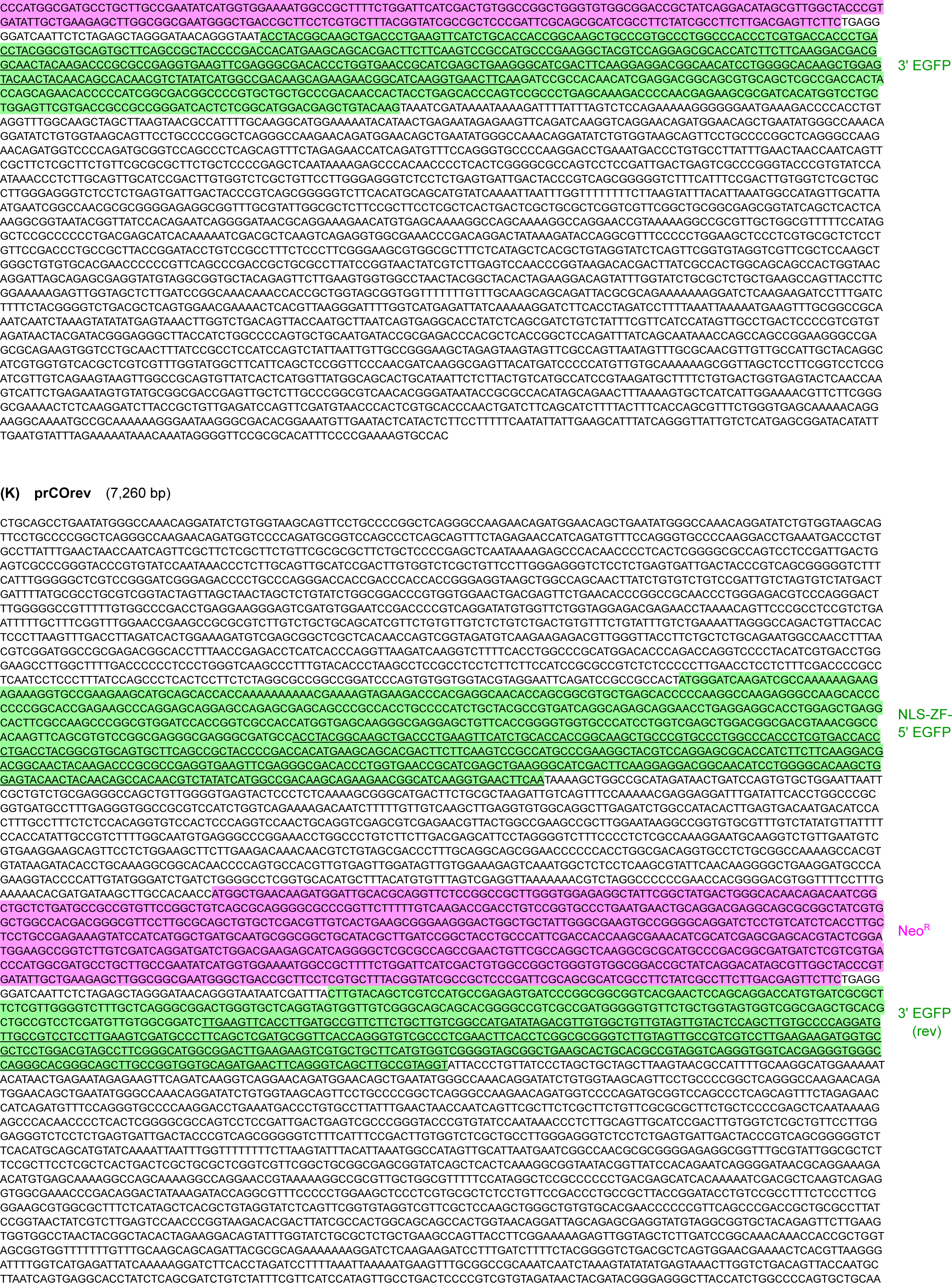
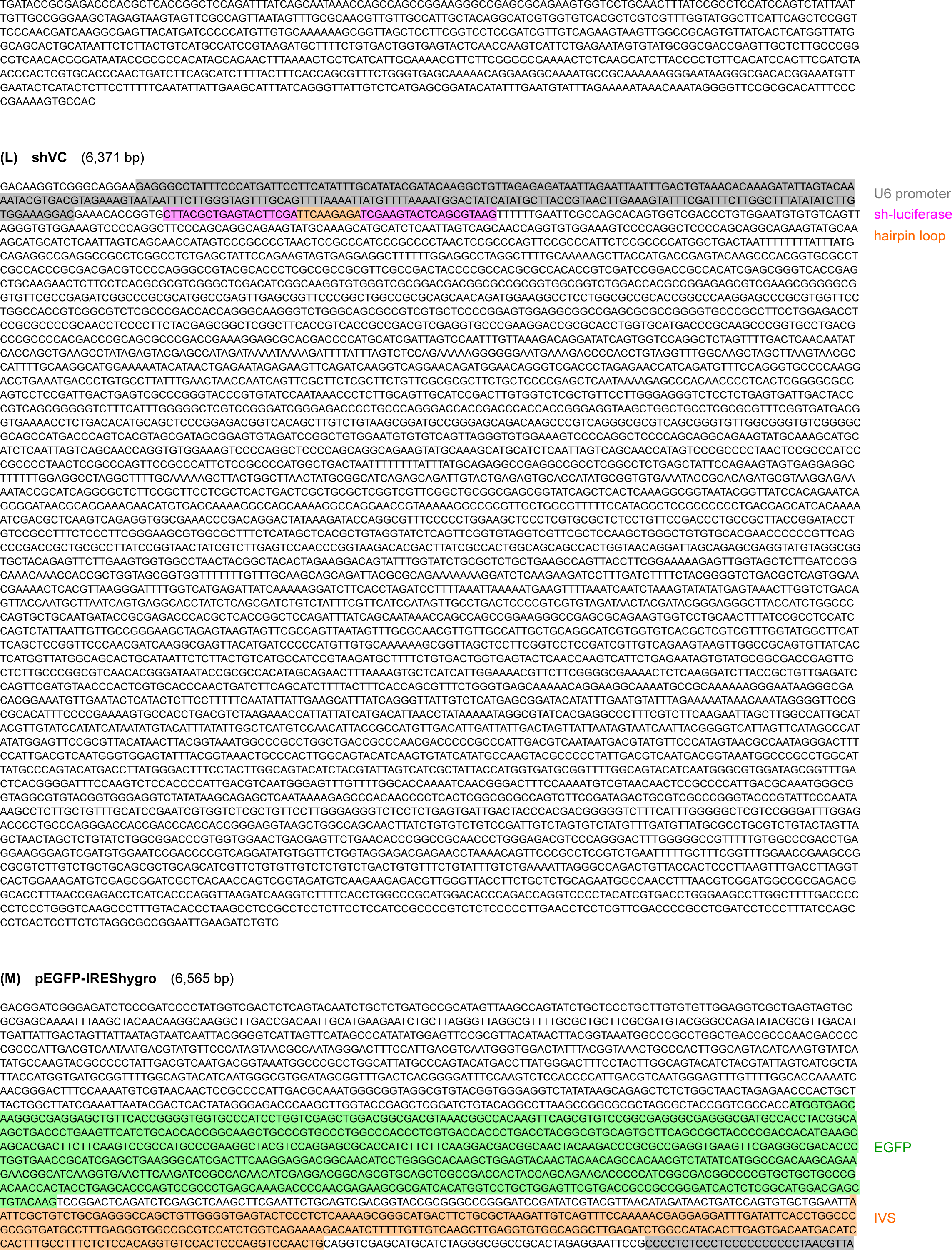
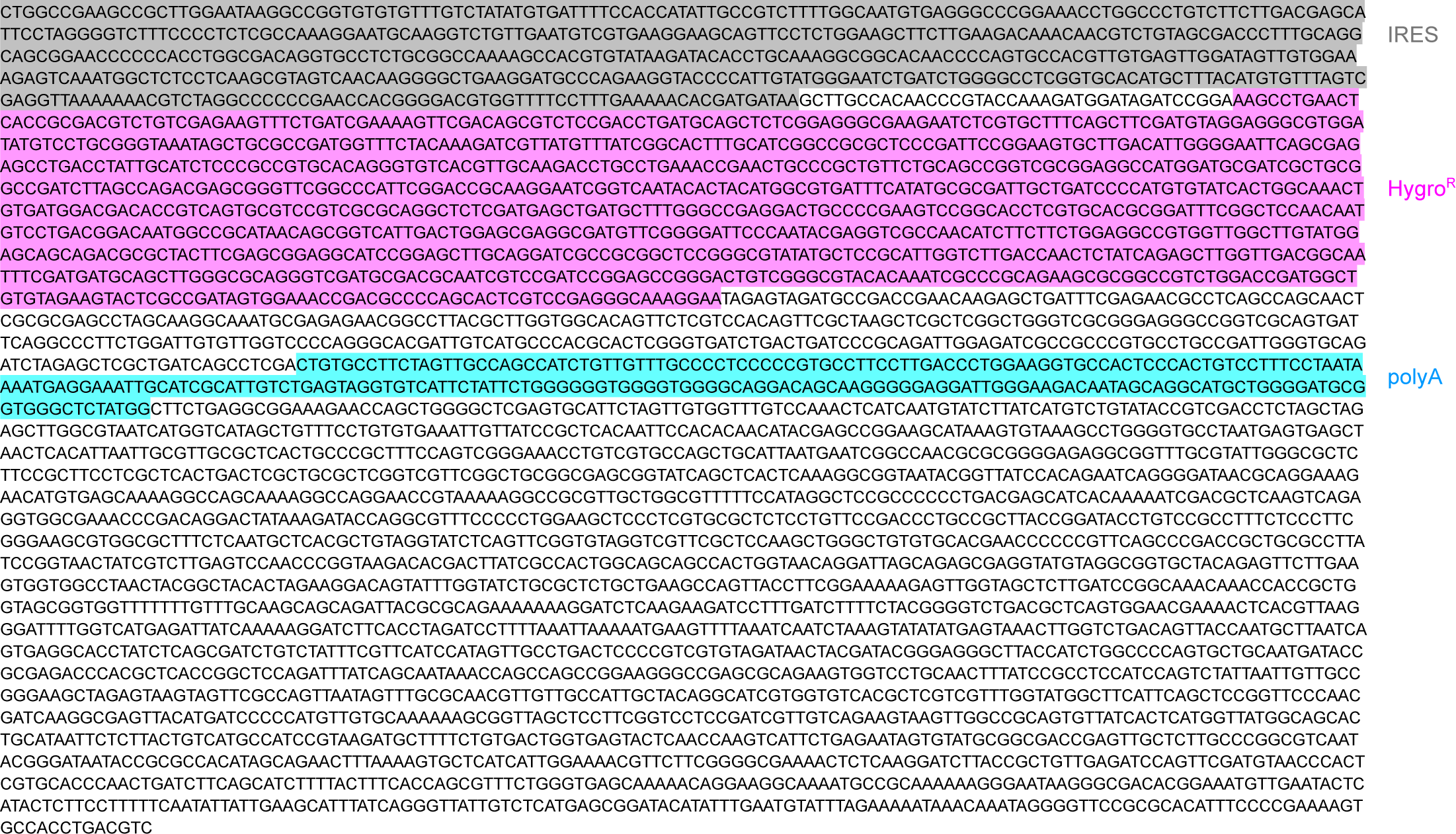
Plasmid sequences. Sequences and features of the plasmids constructed in this study. Donor-*PIGA* (**A**); Donor-*PIGA*^short^ (**B**); Donor-*PIGA*^mut^ (**C**); a vector control for Donor-*PIGA*, Donor-*PIGA*^short^, and Donor-*PIGA*^mut^ (**D**); Donor-*PIGA*-Neo^R^ (**E**); Donor-*CD55*-Neo^R^ (**F**); a vector control for Donor-*PIGA*-Neo^R^ and Donor-*CD55*-Neo^R^ (**G**); pCAG-EGxxFP-*PIGA* (**H**); pCAG-EGxxFP-*CD55* (**I**); prCO (**J**); prCOrev (**K**); shVC (**L**); and pEGFP-IREShygro (**M**). Key features of the plasmids are shaded in different colors and noted on the right. Mutated nucleotides and their wild-type counterparts in the *PIGA* and *CD55* genes are shown in red and green bold letters, respectively. Underlines in (**H**)–(**K**) indicate intramolecular homology between two partial EGFP fragments. PX458-, PX459-, PX461-, PX462-, and pH840Apuro-based Cas9 constructs created in this study are not included in this figure.

**Table S1.**
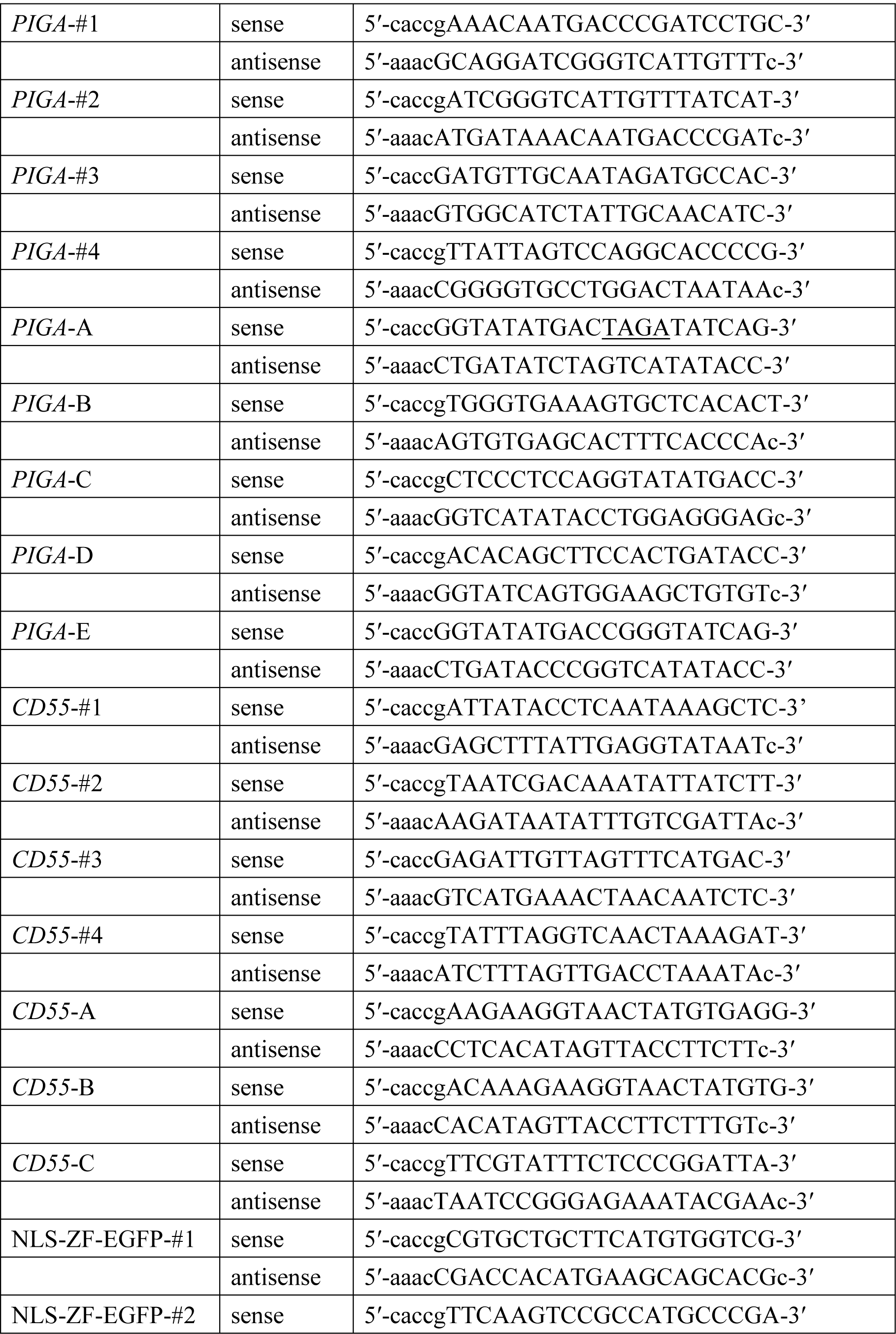
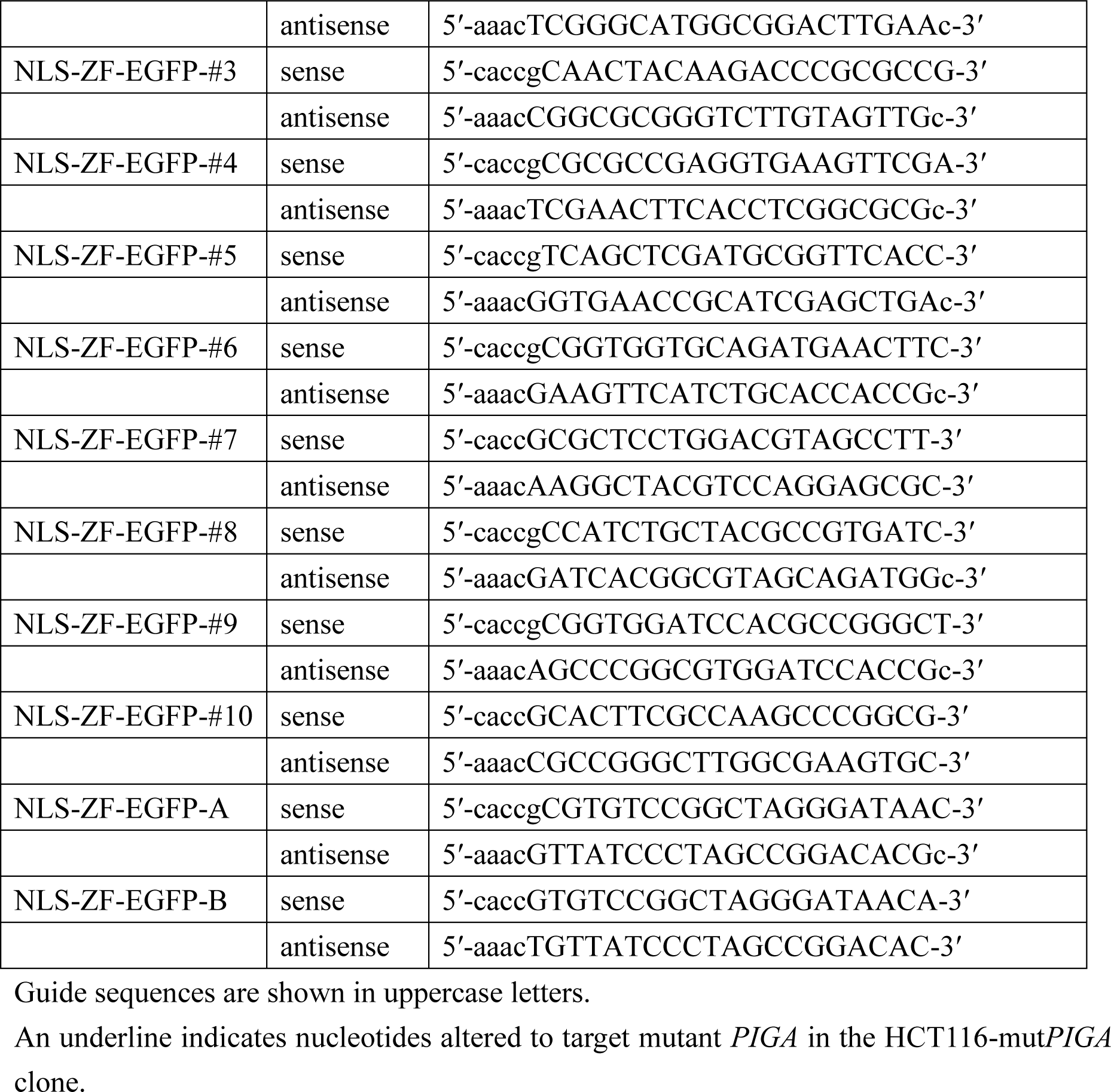
Oligonucleotides used to create sgRNA-expressing vectors.

**Table S2.**
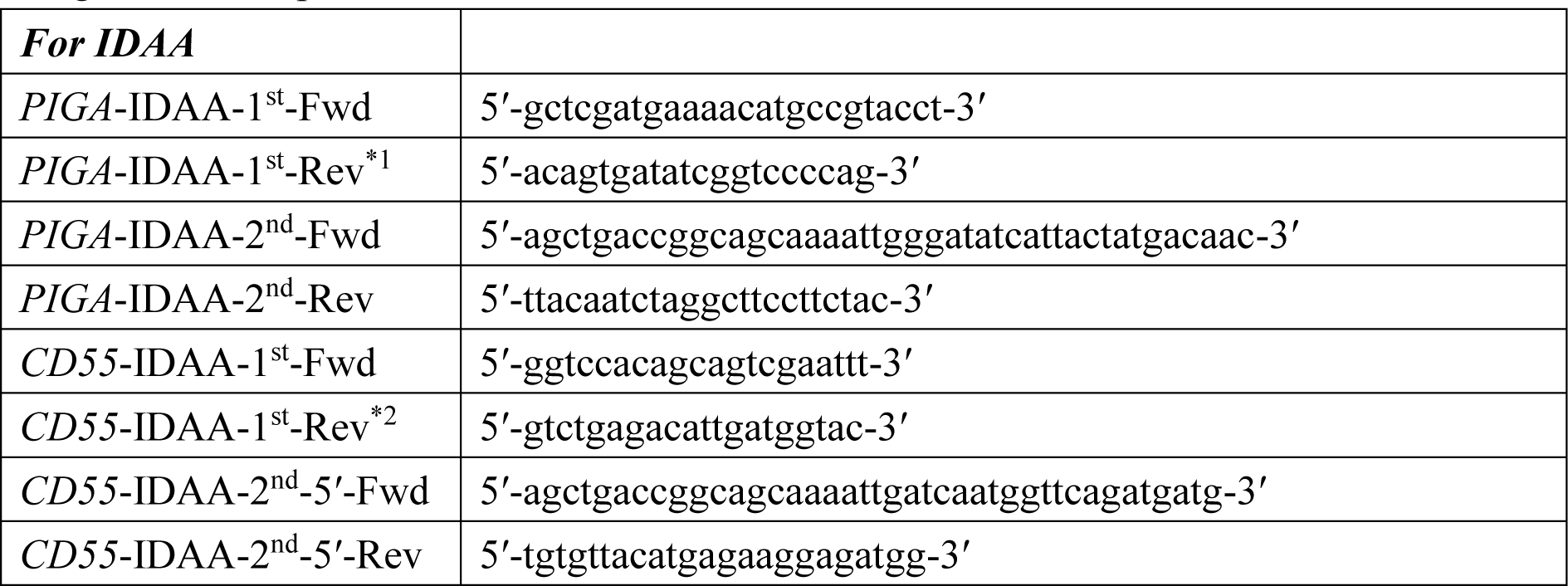
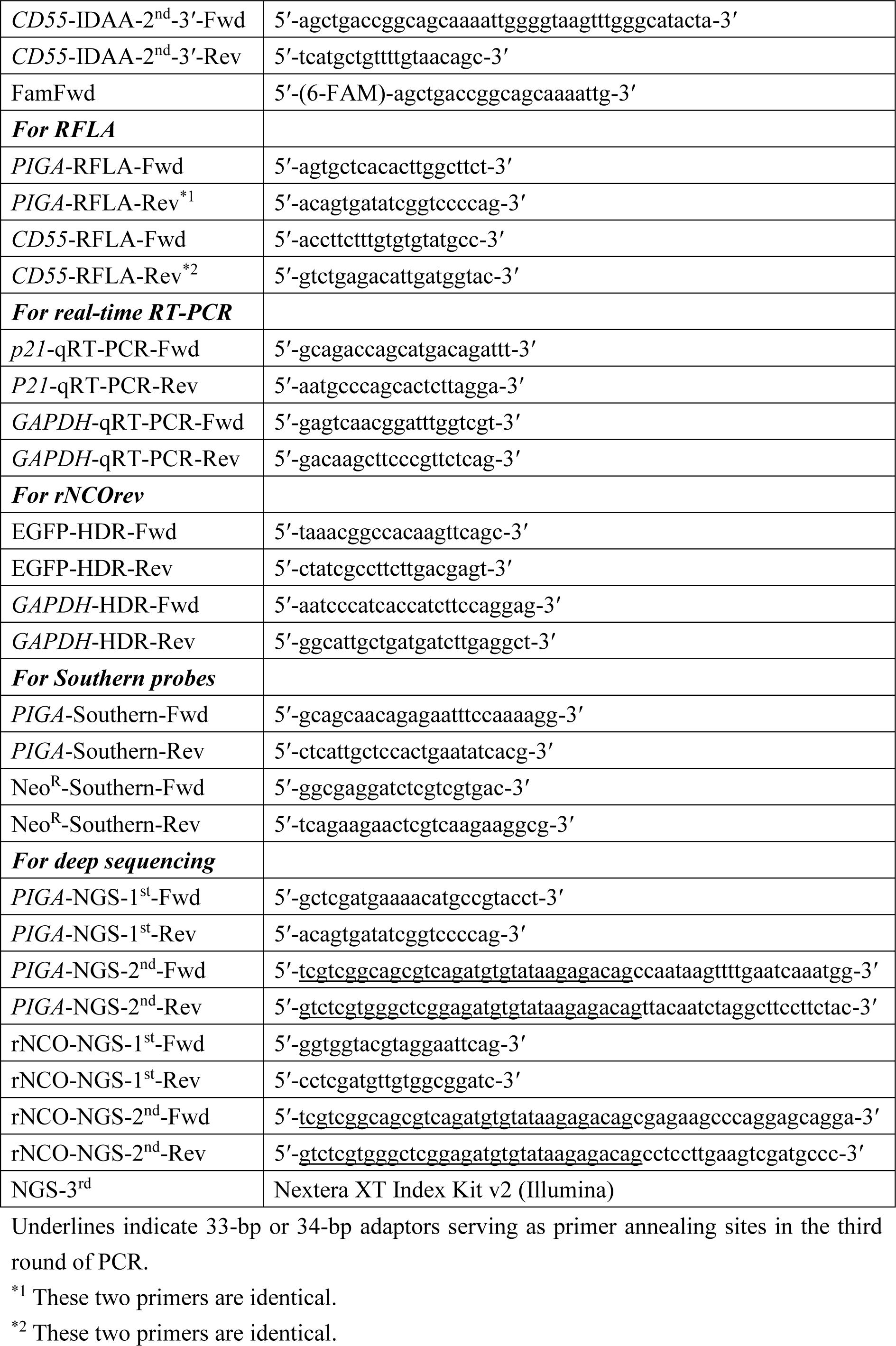
Oligonucleotide primers used for PCR.

**Table S3:**
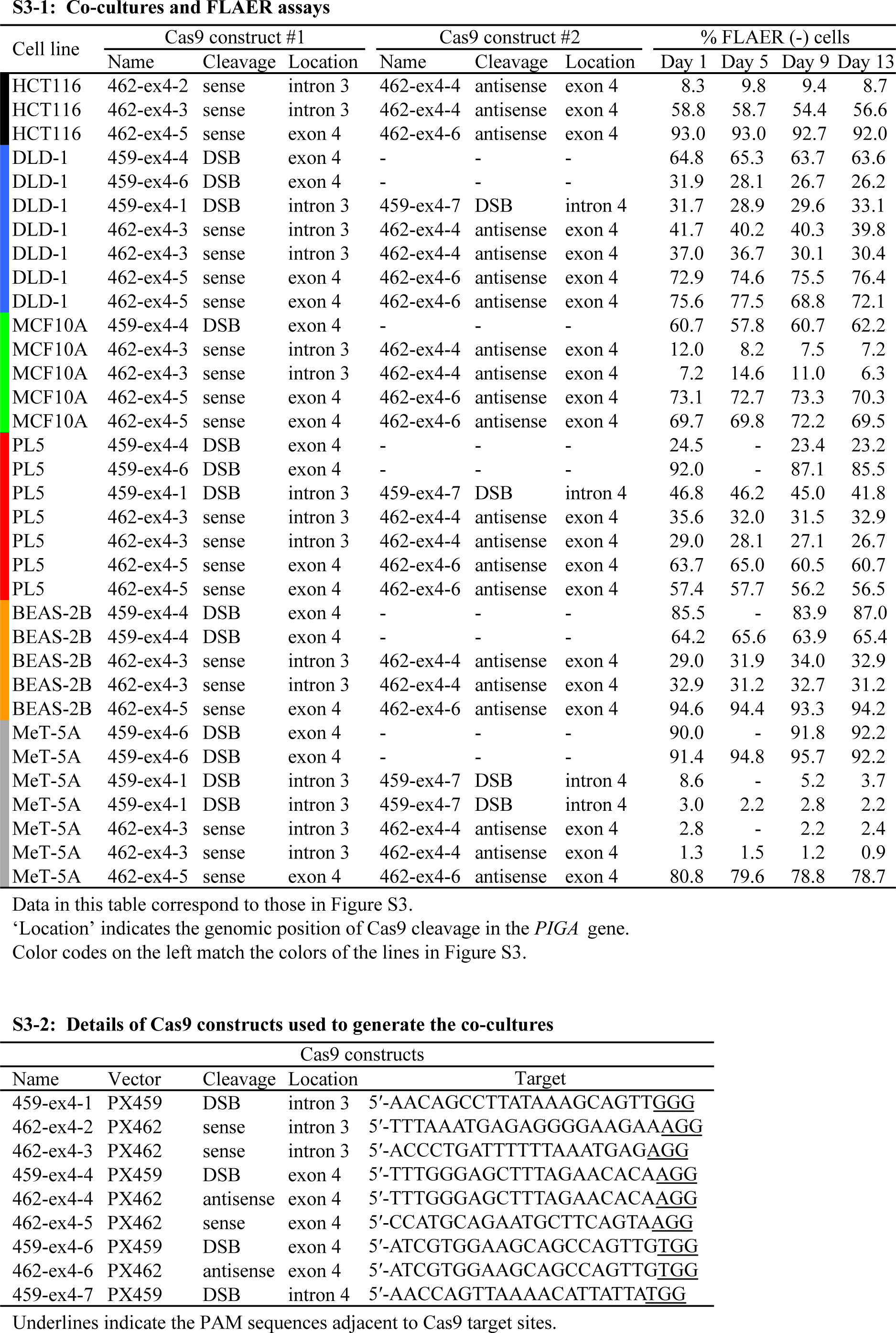
Co-cultures of *PIGA* -positive and -negative cells and FLAER assays.

**Table S4:**
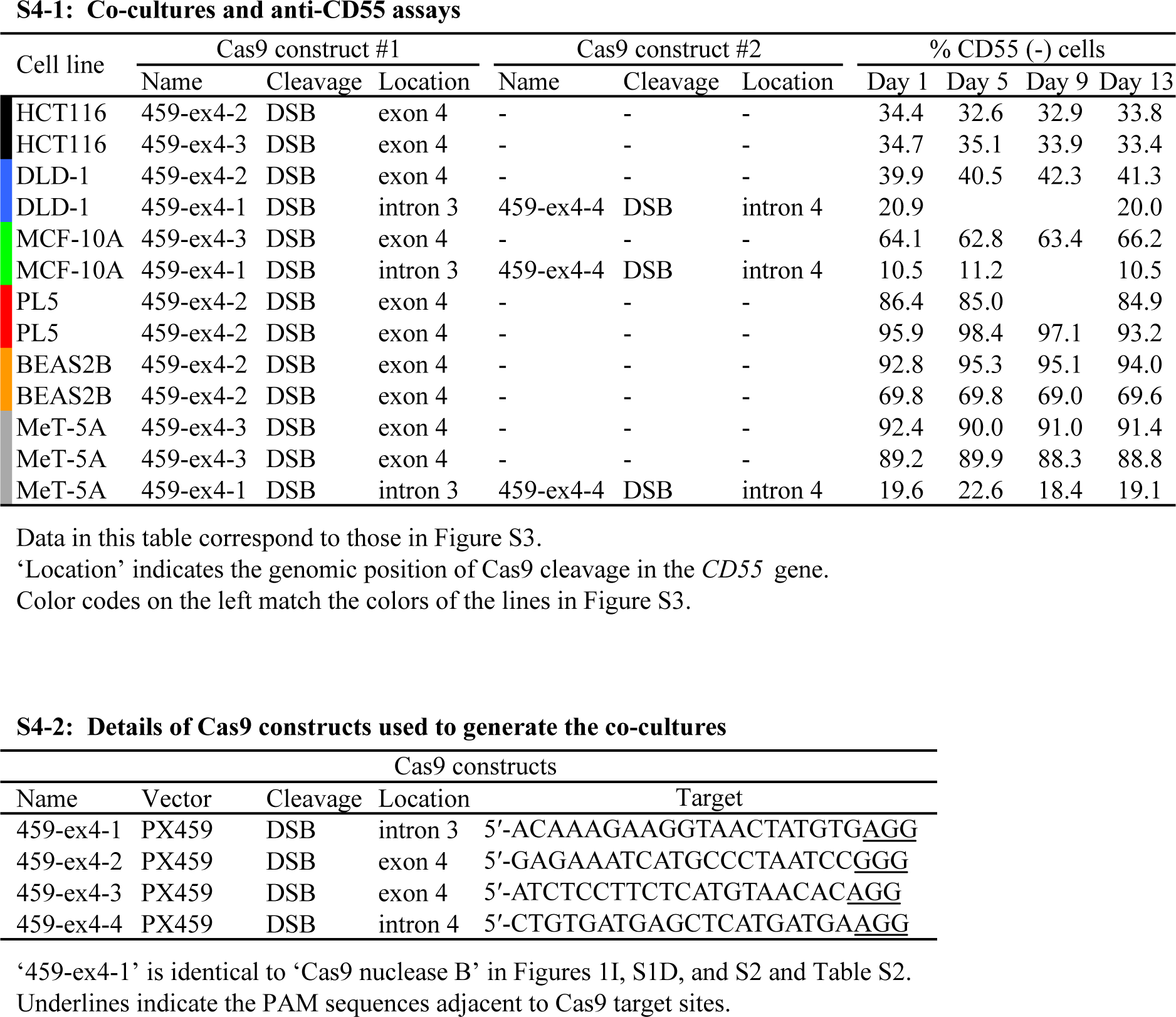
Co-cultures of CD55 -positive and -negative cells and anti-CD55 assays.

